# Functional parcellation of the neonatal brain

**DOI:** 10.1101/2023.11.10.566629

**Authors:** Michael J. Myers, Alyssa K. Labonte, Evan M. Gordon, Timothy O. Laumann, Jiaxin Cindy Tu, Muriah D. Wheelock, Ashley N. Nielsen, Rebecca Schwarzlose, M. Catalina Camacho, Barbara B. Warner, Nandini Raghuraman, Joan L. Luby, Deanna M. Barch, Damien A. Fair, Steven E. Petersen, Cynthia E. Rogers, Christopher D. Smyser, Chad M. Sylvester

## Abstract

The cerebral cortex is organized into distinct but interconnected cortical areas, which can be defined by abrupt differences in patterns of resting state functional connectivity (FC) across the cortical surface. Such parcellations of the cortex have been derived in adults and older infants, but there is no widely used surface parcellation available for the neonatal brain. Here, we first demonstrate that adult- and older infant-derived parcels are a poor fit with neonatal data, emphasizing the need for neonatal-specific parcels. We next derive a set of 283 cortical surface parcels from a sample of n=261 neonates. These parcels have highly homogenous FC patterns and are validated using three external neonatal datasets. The Infomap algorithm is used to assign functional network identities to each parcel, and derived networks are consistent with prior work in neonates. The proposed parcellation may represent neonatal cortical areas and provides a powerful tool for neonatal neuroimaging studies.

**HIGHLIGHTS:**

- Neonatal cortical surface parcels derived based on abrupt changes in functional connectivity (FC) were highly homogenous and were validated in external neonatal datasets.
- Borders between cortical parcels were smoother (less abrupt) in group-average neonatal data compared to adults, likely due to increased heterogeneity in boundary location across individual neonates.
- Parcels derived from adults and older infants show poor fit with neonatal resting-state FC data, underscoring the need for a neonatal-specific parcellation.

## INTRODUCTION

The cerebral cortex is composed of discrete yet interconnected cortical areas that are fundamental macroscale units of the central nervous system. Cortical areas can be defined as contiguous portions of cortex which are distinguished from their neighbors by abrupt changes in function, architectonics, connectivity, and topography (FACT) ^1,2^. Sets of highly interconnected cortical areas, in turn, comprise functional networks that support different aspects of cognition and constitute a higher level of brain organization ^3^. An important goal of human neuroscience is to identify and characterize these cortical areas and functional networks.

‘Parcels’ are subdivisions of the cortex derived empirically in neuroimaging studies based on abrupt transitions in patterns of functional connectivity (FC) across the cortical surface ^4–6^. These abrupt transitions in FC may reflect differences between adjacent cortical areas in function and connectivity ^6^, two of the four FACT criteria, suggesting that parcels could represent cortical areas. Progress has been made in utilizing this method to generate sets of parcels that cover the cortical surface (‘parcellations’) in older infants and adults ^6–9^, and studies using these parcellations have generated a wealth of knowledge regarding adult human brain architecture, function, and relations to individual differences in behavior ^5,10–14^.

Despite the developmental, ontogenetic, and clinical significance of neonatal brain organization, neonatal cortical areas have not been systematically characterized. To date, there are no standard neonatal cortical surface parcellations based on transitions in FC as there are for older infants and adults. While other methods have been used to divide the brain into functionally relevant subdivisions ^15^, these approaches have relied on volume-based rather than surface-based analyses and thus their relations to cortical areas are unclear. Further, adult parcellations are unlikely to fit neonatal data because of the non-linear and non-uniform cortical expansion that takes place over development ^16,17^.

The lack of knowledge of neonatal cortical areas is a significant gap, as the neonatal period is a landmark stage in neural development that serves as a starting point for postnatal experience-dependent learning ^18^. A characterization of the number, locations, and network assignments of neonatal cortical areas is needed to advance our understanding of typical and atypical brain development. A neonatal surface parcellation would also provide a valuable tool for neonatal neuroimaging studies, enabling researchers to reduce dimensionality, increase power by averaging signals over homogeneous regions of cortex, reduce problems of multiple comparisons, and increase methodological consistency across studies and research groups ^19^.

Part of characterizing cortical areas in neonates includes a description of their functional network organization. Functional networks represent a higher level of brain organization and can be defined based on sets of highly interconnected cortical areas or parcels ^3^. Because of the weak long-range anterior-posterior FC of neonates, most prior work describes neonatal networks as anatomically isolated chunks of cortex rather than the distributed organization characteristic of adults ^20–32^. An important goal of developmental systems neuroscience is to characterize the network ‘identities’ of individual neonatal parcels and track the evolving network relations of these parcels over development. Such a description would inform the evolving function of cortical areas over development and provide a foundation for studies of typical and atypical development.

In the current study, we derived a set of neonatal cortical surface parcels based on abrupt differences in FC across the cortical surface in a sample of n=261 neonates. To test the reliability of the parcellation, we split neonates with the most data (primary generation dataset; n=131) into two halves, generated parcellations from each half, and then tested each parcellation on the held-out half of the data. To ensure the generalizability of our parcellation, we then validated it in three external datasets. Finally, we clustered the derived parcels into neonatal functional brain networks. Results provide a robust surface-based neonatal parcellation, uncover putative neonatal cortical areas, reveal properties of neonatal brain network organization, and have practical utility for neonatal neuroimaging studies. The derived parcellation and the code used to derive the parcels is publicly available for use at https://github.com/myersm0/myers-labonte_parcellation/.

## METHODS

This study was approved by the Human Research Protection Office at Washington University in St. Louis. All mothers of neonatal participants provided informed consent prior to study initiation. The primary dataset in this study, eLABE (Early Life Adversity and Biological Embedding), has been recently described ^20,33,34^. The current study focused on fMRI data collected from 261 healthy, full-term neonates (average postmenstrual age, PMA, 41.3 weeks, range 38-45; **Table 1**) from the eLABE dataset scanned between September 2017 and March 2020.

**Table 1:** Full primary dataset demographics table.

| **Neonatal Characteristics (n=261)** | ***n*** | **Mean** | **SD** |
| --- | --- | --- | --- |
| Sex |  |  |  |
| Male | 141 |  |  |
| Female | 120 |  |  |
| Gestational age at birth (weeks) |  | 38.5 | 1.0 |
| Postmenstrual age at scan (weeks) |  | 41.3 | 1.3 |
| Birthweight in grams |  | 3274.0 | 489.9 |
| Area deprivation index |  | 67.9 | 24.9 |
| Child's race |  |  |  |
| African American | 158 |  |  |
| White | 101 |  |  |
| Chinese | 3 |  |  |
| Other Pacific Islander | 1 |  |  |
| Other | 1 |  |  |
| Mixed African American/White | 0 |  |  |
| Mixed Chinese/White | 0 |  |  |
| Ethnicity |  |  |  |
| Hispanic | 6 |  |  |
| Non-Hispanic | 253 |  |  |
| Unspecified | 2 |  |  |
| **Neonatal fMRI Characteristics** | ***n*** | **Mean** | **SD** |
| Amount of fMRI data (minutes) |  | 19.1 | 5.2 |
| Percent of frames retained |  | 16.6 | 4.4 |

### Primary Dataset

Neuroimaging was performed in full-term neonates during natural sleep using a Siemens 3T PRISMA scanner and 64-channel infant specific head coil. Prior to scanning, neonates were fed, swaddled, and positioned in a head-stabilizing vacuum fix wrap ^35^. A T2-weighted image (sagittal, 208 slices, 0.8-mm isotropic resolution, TE = 563 ms, tissue T2 = 160 ms, TR = 3,200 ms) was collected. Functional imaging (fMRI) was performed using a BOLD gradient-recalled echo-planar multiband (MB) sequence (72 slices, 2.0-mm isotropic resolution, TE = 37 ms, TR = 800 ms, MB factor = 8). Using the same parameters, spin-echo field maps were also obtained. Depending on tolerability of the scan, between 2 and 9 fMRI BOLD scans were acquired for each neonate (mean 3.75 runs). Runs were 420 frames, approximately 5.6 minutes in length, and were collected in both the anterior-to-posterior (AP) and posterior-to-anterior (PA) direction; a typical scan session included 2 AP runs and 2 PA runs. AP and PA scans were concatenated following fMRI preprocessing, but prior to FC processing. Framewise integrated real-time MRI monitoring (FIRMM) was used during scanning to monitor real-time neonate movement^36,37^.

### Validation Datasets

Several independent datasets were used to validate results. Each dataset comes from studies that were approved by the Human Research Protection Office at Washington University in St. Louis. The first validation set, CUDDEL+OXYGEN, is a combination of two datasets, Prenatal Cannabis Use and Development of Offspring Brain and Behavior During Early Life (CUDDEL) and Maternal Oxygen in Labor (OXYGEN), both identical to the primary dataset (eLABE) in recruitment methods, demographics, neuroimaging acquisition, and processing. CUDDEL is enriched for mothers who reported cannabis use during pregnancy, and 33% of the cohort used in these analyses were exposed to cannabis in utero. OXYGEN recruited mothers during labor with the goal of scanning healthy neonates within the first 72-hours of life. As both CUDDEL and OXYGEN were ongoing studies during the time of our analyses, we restricted our use of the data to the sample sizes that were available at the start of these analyses (n = 36 and n = 5, respectively). **Supplementary Figure 1** shows that we obtained robust results with these sample sizes.

A second validation dataset, Precision Baby (PB003), consists of a single neonate (Male; 44-weeks PMA) scanned across 5 consecutive days for a total of 3.27 hours of low-motion fMRI data (FD < 0.25). Recruitment of this neonate was the same as the primary dataset (eLABE), and neuroimaging acquisition and processing were also largely similar. A T2-weighted was collected using the same parameters as the primary dataset and was used to align functional data across all scan sessions for each neonate. Functional imaging (fMRI) was performed using a BOLD gradient-recalled echo-planar multiband sequence (60 slices, 2.4-mm isotropic resolution, TE = 37 ms, TR = 1200 ms, MB factor = 4). Using the same parameters, spin-echo field maps were also obtained.

The third external validation dataset, WUNDER, consists of 70 full-term neonates scanned between 2007 and 2016, and has been previously described ^24^. Neuroimaging for this dataset was performed during natural sleep using a Siemens 3T TRIO scanner and infant-specific head coil. Structural images were collected with a turbo spin-echo T2-weighted sequence (TE=161 ms, TR=8950 ms, 1 mm isotropic resolution). Functional imaging (fMRI) was performed using a gradient-echo echo planar image sequence sensitized to T2* BOLD signal changes (2.4 mm isotropic resolution, TE = 28 ms, TR = 2910 ms). Spin-echo field maps were also obtained using the same parameters. A total of 200 frames were collected over 10 minutes for each run. A minimum of one run was required for inclusion, but some infants had up to four runs.

### fMRI Preprocessing

All datasets underwent identical preprocessing and FC processing except WUNDER, which is described below. fMRI preprocessing included correction of intensity differences attributable to interleaved acquisition, linear realignment within and across runs to compensate for rigid body motion, bias field correction, intensity normalization of each run to a whole-brain mode value of 1,000, distortion correction, and linear registration of BOLD images to the adult Talairach isotropic atlas ^38^ using in-house software (ftp://imaging.wustl.edu/pub/raichlab/4dfp_tools/). Field distortion correction was performed using the FSL TOPUP toolbox (http://fsl.fmrib.ox.ac.uk/fsl/fslwiki/TOPUP). BOLD images for each subject were first registered to their individual T2 and then to a cohort-specific T2 atlas in 711-2N Talairach space. The cohort-specific T2 atlas was created from a subset of 50 neonates from the eLABE dataset.

The volumetric preprocessed BOLD data were then mapped to subject-specific surfaces prior to FC-processing. The Melbourne Children’s Regional Brain Atlases (MCRIB) ^39,40^, a surface-based neonatal tissue segmentation approach, was used to generate surfaces for each subject from their T2 image that had been linearly transformed to adult Talairach (711-2N) space. Subject-specific surfaces were aligned across subjects into the “fsLR_32k” surface space using spherical registration procedures ^41^ adapted from the Human Connectome Project as implemented in Connectome Workbench 1.2.3 ^42,43^. All volumetric and surface registrations were visually inspected to ensure accuracy.

Following initial preprocessing and surface registration to fsLR_32k space with a small smoothing kernel (σ = 1 mm), BOLD time series were censored at framewise displacement (FD) < 0.25 mm, and only epochs of at least 3 consecutive frames with FD < 0.25-mm were included. A minimum of 10 minutes (750 frames) of usable data were required from each subject for inclusion. This data then underwent FC processing as follows ^6,44^: (1) demean and detrend within each run, ignoring censored frames; (2) multiple regression with nuisance time series including white matter, ventricles, and whole brain (average gray matter signal), as well as 24 parameters derived from head motion, ignoring censored frames. Finally, retained data were interpolated at censored timepoints to allow band-pass filtering (0.005 Hz < f < 0.1 Hz).

Time courses for surface data were smoothed with geodesic 2D Gaussian kernels after FC processing (σ = 2.25 mm). FC was computed as the Fisher z-transformed Pearson correlation between time courses from pairs of surface parcels or vertices, as detailed below.

fMRI preprocessing for the WUNDER dataset was similar except that the time series were high pass filtered (<0.08 Hz) and spatially smoothed (σ = 6 mm). Only subjects having at least 5 minutes of low-motion fMRI data were included in the analysis; this more lenient threshold was allowed for the older WUNDER dataset to include an many subjects as possible in the group-average.

### Boundary Map Generation

Following the notion that adjacent cortical areas should be separated by abrupt changes in function and connectivity^1^, the method described in ^6^ was used to identify transitions in FC across the cortical surface. First, pairwise FC of all ∼60k cortical surface vertices was computed for each subject, resulting in a 60k x 60k square RSFC matrix where each row or column is the “correlation map” of a particular vertex: a vector of values characterizing a single vertex’s correlation with all other vertices. Next, a 60k x 60k ‘similarity matrix’ was generated for each subject as the pairwise correlations between all the correlation maps; the similarity matrix thus represents how similar the correlation maps are between every possible pair of vertices. The first spatial derivative of each subject’s similarity matrix was then computed, producing a 60k x 60k matrix in which each column is the ‘gradient map’ of a particular vertex, representing how abruptly that vertex’s similarity in RSFC with other vertices changes as one moves across the cortex. Vertex-wise gradient maps were then averaged across all subjects, creating an across-subjects average gradient map for each vertex, and smoothed with a kernel of σ = 2.55 mm. The watershed algorithm ^45^ was applied to each vertex’s average gradient map. With this method, regions are “filled up” from their local minima (low points in the gradient map) until they reach border vertices that could be assigned to more than one region; such borders represent locations of peak spatial gradient (i.e., rapid changes in FC similarity) across participants. We experimented with a parameter to control the number of bins into which continuous-valued heights are discretized for iteration within the watershed algorithm. In the original algorithm outlined for adults in (Gordon et al. 2016), this steps parameter was set to 400 bins to improve computation time; however, noting that higher values of this parameter (i.e., smaller bins for each iteration of the watershed) produced sharper edge maps in our dataset, we modified this parameter to 1600 bins. This parameter was also modified and set to 1600 to recompute the adult boundary maps from Gordon et al. ^6^ to show fair comparisons between adults and neonates. The binary maps of gradient peaks (borders) were then averaged across all gradient maps to produce a “boundary map”, which represents the probability of each particular vertex being classified as a border. A flowchart of this method can be seen in **Supplementary Figure 2**. Adult and neonatal boundary maps were compared with a smoothness metric using a built-in Connectome Workbench command (wb_command-cifti-estimate-fwhm).

### Parcel Generation

Parcels were generated using the same watershed procedure described above to fill the boundary map from its local minima. Several parameters can influence the sizes and shapes of the resulting parcels. One of these parameters concerns the tendency for two neighboring parcels to be combined into one parcel. Two neighboring parcels may be combined into a single parcel if a relatively small boundary between them suggests that their respective connectivity patterns are not sufficiently different. To select an appropriate value for this parameter, we visually inspected a range of thresholds (from 0.25 to 0.4) and chose the value (0.38) that best respected the salient boundaries we observed in the boundary map.

The second parameter concerns the height at which we stopped “filling” the regions, the height criterion. In the original method outlined for adults in ^6^, this value was selected as the 90^th^ percentile of all height values, such that the 10% of vertices exceeding this height were left unassigned (i.e., not belonging to any parcel). These unassigned values may be considered transitional zones in which the connectivity is rapidly changing. This parameter determines the approximate percentage of the cortex that is to be covered by the parcellation. We tested several thresholds for this height criterion parameter, ranging from the 25^th^ to 90^th^ percentiles (see **Figure 4**). The highest homogeneity of the parcels in external datasets was observed using the 50% height threshold. Thus, this threshold was used to generate the parcellation used for all subsequent analyses.

The ‘final’ parcellation was generated using the procedure above in the primary dataset, the half of subjects with the most data following frame censoring (n=131). This restriction ensured that the parcellation was generated from subjects with data with the lowest noise, as well as to test the parcellation in the held-out half of the as described in the results (‘Parcellation generated from neonatal data has high homogeneity in external datasets’). This procedure resulted in 304 parcels tiling the cortical surface, 153 parcels in the left hemisphere and 151 parcels in the right. Parcels with fewer than 15 vertices outside of the low signal areas (mean signal <750 after mode 1000 normalization) ^6,46^ were removed because they were unlikely to be reliable. This procedure removed 22 parcels primarily in the inferior temporal and orbitofrontal lobes, resulting in 282 parcels tiling the cortical surface. We also identified four parcels that did not appear biologically plausible because of their shapes. Parcel #217 was trimmed. One parcel with a biologically implausible shape was split into two parcels (#4, #137). We removed one vertex from parcel #33. We added two vertices to parcel #121 to fill in a hole in the middle. The resulting final parcellation consisted of 283 parcels, with 146 parcels in the left hemisphere and 137 in the right.

### Parcel Homogeneity

The parcel generation procedure outlined above creates parcels based on distinct boundaries which indicate differences in FC patterns among adjacent cortical regions ^6^. These generated parcels should both be distinct from neighboring parcels in connectivity and show homogenous connectivity within each parcel. To measure the homogeneity of a parcel, a principal components analysis (PCA) is run in which the inputs are the connectivity patterns from all the individual vertices comprising a parcel in the group average data. Following ^6^, we define homogeneity as the percent of variance explained by the first component.

Homogeneity is highly related to the size of a parcel such that smaller parcels tend to have higher homogeneity than larger parcels ^6,47^. To provide a fair point of comparison, we considered a null distribution of random parcellations having parcels of the same sizes and shapes, and in the same configuration as our true parcellation, but randomly relocated about the cortical surface. To do this, we replicate the rotation method described in ^6^. Briefly, the original parcellation was randomly rotated around each of the x, y, and z axes on a spherical expansion of the cortical surface, allowing for the random relocation of each parcel while maintaining their size, shape, and relative positions to one another. This rotation procedure was repeated 1000 times, whereby each hemisphere was rotated symmetrically with each iteration. The average homogeneity of the original parcellation (described above) was then compared with the homogeneity values of each of the randomly rotated parcellations, calculated as a z-score ((original homogeneity – mean of random homogeneities)/standard deviation of random homogeneities).

### Parcel Reliability

To test the reliability of our parcellation, we randomly split the primary dataset (n=131) in half, generated parcellations from each half separately, and evaluated the overlap of resulting parcels. To quantify the spatial overlap, the Dice similarity coefficient (DSC) was computed on binarized parcel identity maps for the two halves. To assess the significance of this result, we randomly rotated both parcellations 1000 times (previously described in detail under ‘Parcel Homogeneity’), each time computing the DSC of the two randomly rotated parcellations to derive a null distribution against which to compare the value obtained in the true parcellations.

### Identification of Parcel Network Structure

The community detection algorithm Infomap was used to empirically derive functional brain networks from our parcels ^48,49^. For each subject in the primary dataset, we created a parcellated time series by calculating the mean within-parcel time series over each of the 283 parcels. We then cross-correlated these parcellated time series to generate a parcel-wise correlation matrix. Parcel-wise correlation matrices were Fisher z-transformed and averaged across all subjects to generate a single parcel-wise correlation matrix, which was then masked to remove functional connections with a distance < 30mm along the cortex between the nearest vertices of each pair of connected parcels. Functional connections surviving this distance threshold were then binarized to isolate only the strongest positive connections over a range of thresholds chosen to achieve varying degrees of sparseness (in 40 steps ranging from 0.25% to 10%). The resulting 40 connection matrices were then used separately as inputs to the Infomap algorithm, to assign parcels to communities (or networks) based on the maximization of within-community random walks in the connection matrix. This produced 40 network solutions, one for each of the edge densities.

Putative network identities were then assigned by matching communities at each threshold to a set of previously described neonate-specific vertex-wise networks (**Supplementary Figure 3**) (from the 1.25% edge density vertex-wise Infomap solution as illustrated in Figure 3A in ^20^). This matching approach proceeded as follows. At each density threshold, all identified communities were compared (using spatial overlap, quantified with the Jaccard index) to each of the networks in turn. The best-matching (highest-overlap) parcel-wise community was assigned that network identity; that community was not considered for comparison with other networks within that threshold. Matches lower than Jaccard = 0.1 were not considered (to avoid matching based on only a few vertices). Matches were first made with larger networks (Anterior and Posterior Default, Lateral Visual, Motor, Fronto-Parietal, Dorsal Attention), and then to the smaller networks (Orbito-Frontal and Premotor). A “consensus” network assignment was then derived by collapsing assignments across thresholds, giving each parcel the assignment it had at the sparsest possible threshold by which it was successfully assigned to one of the known networks.

A few manual modifications were made to the “consensus” network assignments, which were evident in Infomap solutions, but that did not appear in the neonate-specific vertex-wise network template used in the consensus procedure described above. The motor network, which appeared as a single network in the template, was divided into motor hand and motor mouth networks, based on assignments at the sparest edge threshold (0.25%). Further, parcel number 161 was changed from the lateral visual to the medial visual network, based on the network that had been assigned at sparser edge densities rather than the assignment from the consensus algorithm described above. This decision was made based on the overall pattern across edge densities. Brain surface visualizations were generated in Julia version 1.8.3 with the Makie plotting library ^50^.

Spring embedded plots were also generated to visualize how clustering patterns of parcels changed across various edge densities. All plots were generated using the igraph library in R ^51^.

## RESULTS

### Performance of Adult and Infant Parcels on Neonatal Data

We first investigated how well parcels generated from adults and older infants perform on neonatal data to determine the need for a neonate-specific areal parcellation. Publicly available adult (Glasser et al. 2016; Schaefer et al. 2018; Gordon et al. 2016) and infant parcellations ^9,15,52^ were tested on the half of the dataset which was not used to generate the neonatal parcellation (n=130). Most of the adult parcellations performed no better than chance in neonatal data (Schaefer: p=0.771, Glasser: p=0.012, Gordon: p=0.658) (**Supplementary Figure 4**). While the Glasser et al. parcellation performed slightly better than chance, the z-score of 2.3 was much lower than z-scores typically reported for well-fitting parcellations ^6^. Parcellations created from older infants similarly did not perform better than chance in our neonatal data (Wang 0-2Yr: p=0.188, Wang 0-3Mo: p=0.247, Shen: p=0.932) (**Supplementary Figure 4**). Thus, parcellations specific to adults and older infants do not capture neonatal FC patterns, underscoring the need for neonate-specific parcellations.

### Neonatal Boundary Map

A neonatal boundary map was generated from the primary dataset (n=131; **Supplementary Table 1**), identifying transitions in patterns of FC across the cortex (**Figure 1)**. Neonatal boundaries (**Figure 1A**) appeared thicker and smoother compared to an adult dataset (**Figure 1B**). Quantitatively, the spatial smoothness of the neonatal boundary map was 4.06 mm FWHM, compared to 3.2 mm FWHM for the adult group average boundary map, suggesting that the borders are less ‘sharp’ in data averaged across neonates compared to data averaged across adults. In addition, we noted that smoothness of the neonatal boundary map increased with increasing number of neonates included in creating the average (**Supplementary Figure 5**), suggesting that the smoothness of borders may result from variability in border placement across neonates. Consistent with this hypothesis, the spatial smoothness of a boundary map from an individual neonate (PB003) was 2.82 mm FWHM, comparable to the adult group average boundary map (**Figure 1C**).

**Figure 1:**
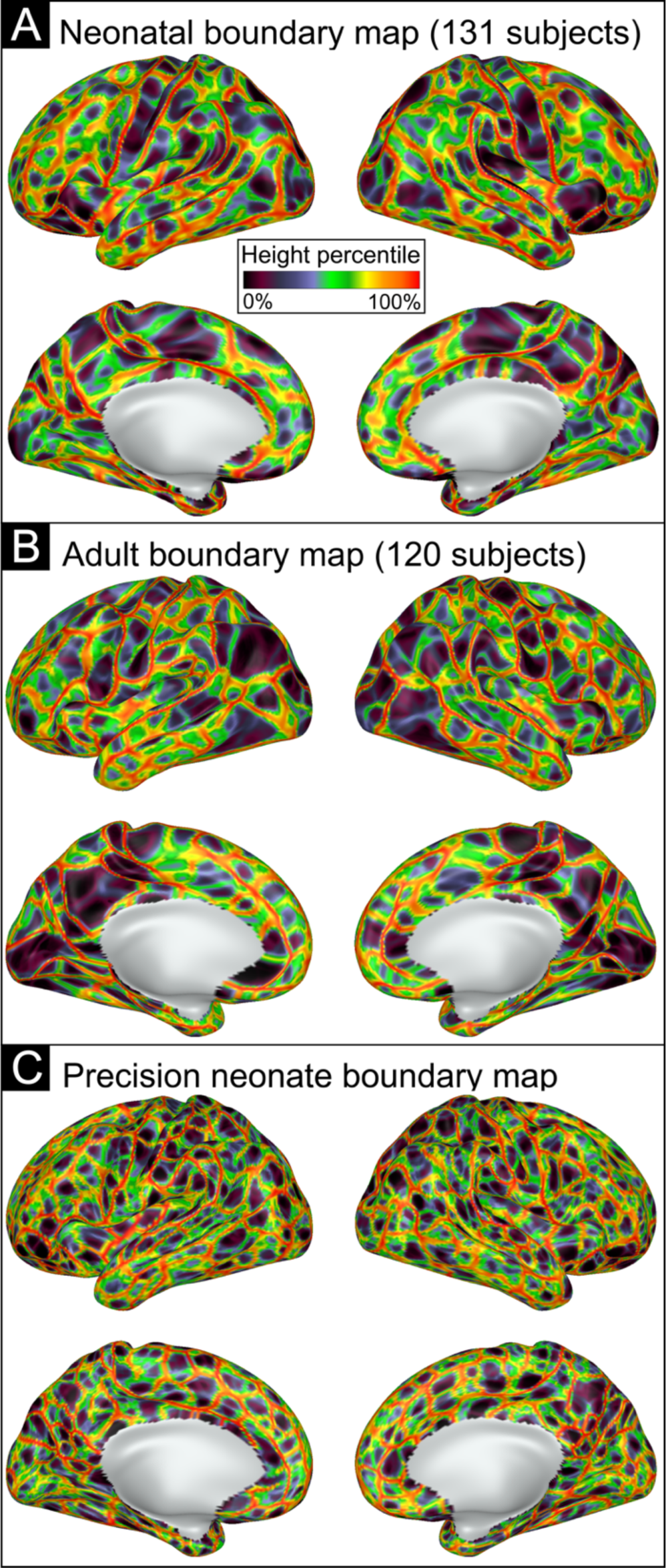
Resting state functional connectivity (RSFC) boundary maps generated based on abrupt changes in FC. A) Neonatal RSFC boundary map generated based on the average of all 131 subjects’ gradient maps. B) Adult RSFC boundary map based on the average of 120 subjects. C) Neonatal RSFC boundary map generated based on a single neonate’s gradient maps. Boundaries are indicated by color based on height percentile of edge density, where bright colors indicate locations where abrupt transitions in RSFC patterns were consistent across many cortical vertices, and darker colors represent areas of cortex where the RSFC patterns were relatively stable.

### Neonatal Parcel Creation

The primary dataset (top half of participant in terms of data quality; n=131) was split randomly into split-half 1 (n=65) and split-half 2 (n=66). **Supplementary Table 2** shows demographic information for each split half. For each split half, we created a separate boundary map (similar to Figure 1A) and then generated a series of parcellations over a range of height criteria (from the 25^th^ to the 90^th^ percentile of height values). **Figure 2A** illustrates how each split-half parcellation performed at each height threshold against both its generating sample and the held-out sample. Parcellations performed extremely well when tested against the held-out sample (z ∼9-10) at lower height thresholds (25^th^ through 50^th^ percentiles). Performance steadily declined as height criteria increased beyond an inflection point at the 50^th^ percentile. Performance was lowest at the 90^th^ percentile but still significantly better than chance for the held-out samples (z close to or above 3.3, p<0.001). Thus, we concluded that the optimal height criteria for our neonatal parcellation in terms of generalizability and within-sample testing was 50%.

**Figure 2:**
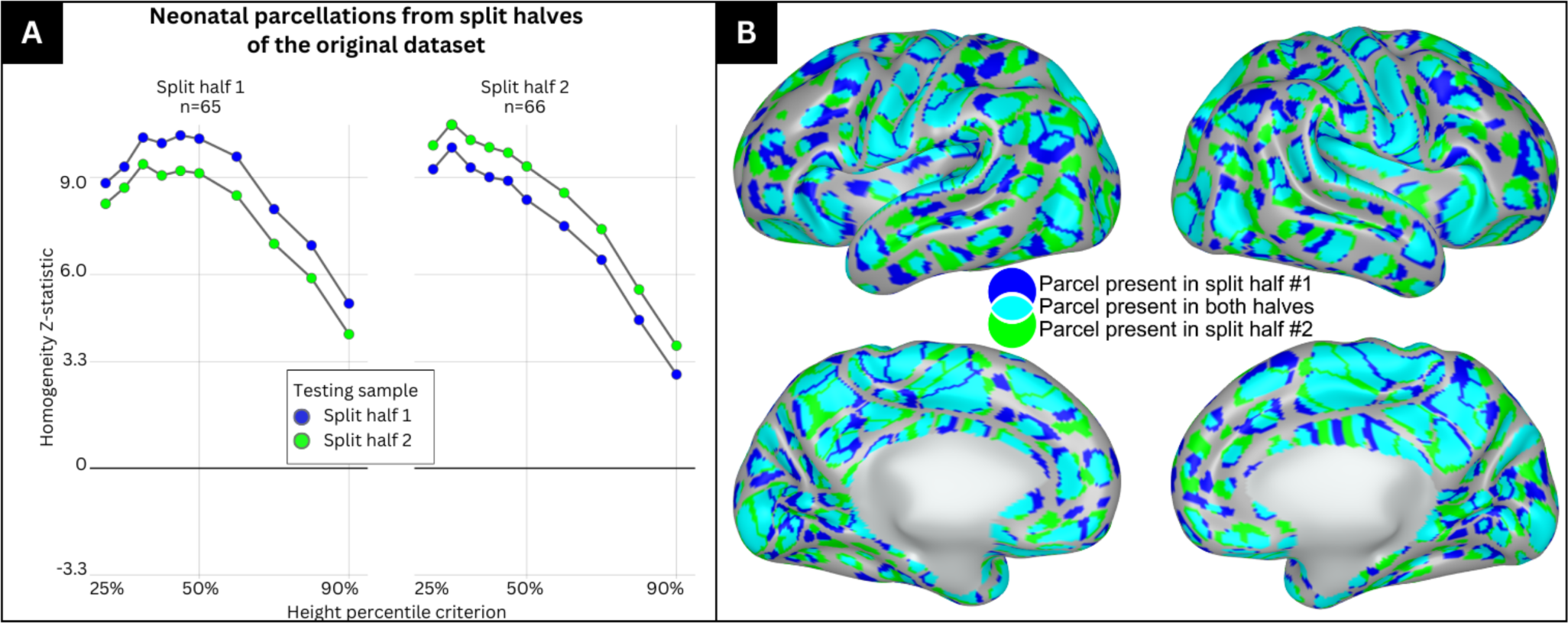
Parcels generated at the 50% height threshold from split-halves of the primary dataset highly overlap with one another. **A)** The primary dataset was split into split half 1 and split half 2 and used to generate parcellations at varying height thresholds between 25% and 90%. Each split-half parcellation was then tested against the sample that generated the parcellation and the other split half. The left panel represents the homogeneity z-statistic of the parcellation generated from split half 1 tested against itself (blue) and the other split half (green) at varying height thresholds. The right panel represents the homogeneity z-statistic of the parcellation generated from split half 2 tested against itself (green) and the other split half (blue) at varying height thresholds. **B)** Medial and lateral view of the right and left hemisphere showing parcels which are identified in split half 1 only (blue), split half 2 only (green) and in both split halves (cyan).

Figure 2B illustrates the parcels generated from each split-half at the 50% height threshold. Parcels from split-half 1 are shown in blue, parcels from split-half 2 are shown in green, and areas of parcel overlap are colored cyan. Visual inspection of Figure 2B indicates good overlap between the parcels generated from the two split halves across most of the cortical surface. The Dice coefficient of overlap in parcels from the two split halves was 0.69 and was highly significant based on rotation-based null models (z = 19.5).

Our ‘final’ parcellation was generated from the primary dataset (n=131) at the 50% height criterion and is shown in Figure 3. There were 283 parcels in this final parcellation, with 146 parcels in the left hemisphere and 137 in the right.

**Figure 3:**
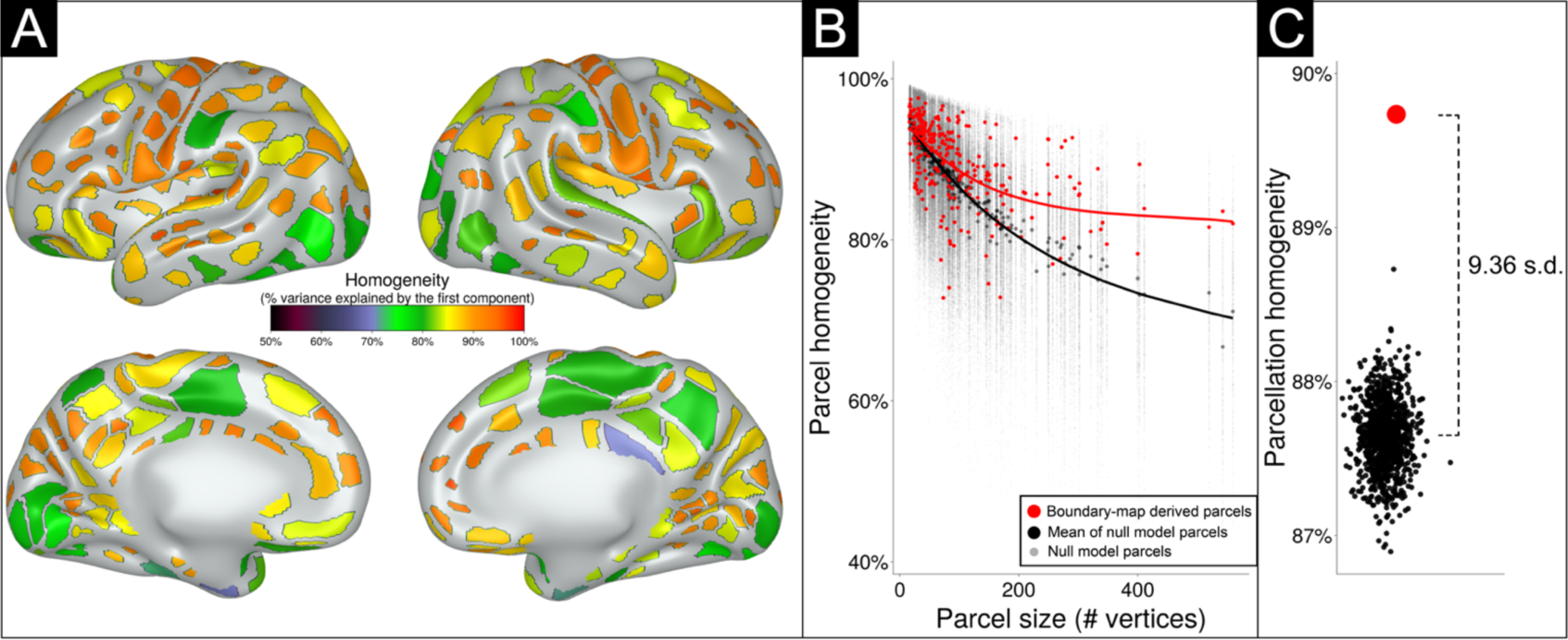
The parcellation generated from the primary dataset (n=131) shows 283 highly homogenous parcels at 50% height threshold. **A)** Parcels are colored based on homogeneity value, calculated based on percent variance explained by the first principal component of the connectivity patterns of the individual vertices comprising each parcel. **B)** The homogeneity of each parcel (red dots) is plotted as a function of parcel size. Black dots indicate the homogeneity of each parcel over 1000 null rotations. Note that many parcels had true homogeneity values higher than the average of null rotations, especially larger parcels. **C)** The performance of the entire parcellation scheme tested against 1000 null rotations. The black dots indicate the average homogeneity value across all 283 parcels for each of the 1000 null rotations. The red dot represents the average homogeneity value across all 283 parcels in the true data. The average homogeneity across all parcels was 9.36 standard deviations above the mean of the null rotations.

### Neonatal parcellation has high homogeneity

The homogeneity value of each parcel in the final parcellation is illustrated in Figure 3A. Most generated parcels (red dots) had homogeneity values higher than expected by chance compared against random parcel rotations (black dots) on the cortical surface (Figure 3B). As depicted in Figure 3C, and as previously noted ^6^, larger parcels tended to be less homogenous; however, larger parcels tended to do better against the null rotations compared to smaller parcels, due to smaller parcels having high homogeneity in many of the null rotations. The parcellation as a whole also had higher homogeneity averaged across all 283 parcels (Figure 3C; red dot) compared against the average homogeneity from any of the 1000 null rotations (p<0.001) (Figure 3C; black dots). The homogeneity of the real parcellation was 9.36 standard deviations above the mean of the null rotations.

### Parcellation generated from neonatal data has high homogeneity in external datasets

The final neonatal parcellation was validated against three separate external neonatal datasets by comparing the average homogeneity across all parcels against average homogeneity from 1000 null rotations. In Figure 4A, bars illustrate testing of the parcels against the excluded half of the dataset (the half of the dataset with lower amounts of retained data after censoring; n=130) and against three different external datasets. The parcellation generated from the primary dataset performed very well against its excluded half (z=9.0), the CUDDEL external group dataset (z=8.92), and the precision dataset (PB003) from a single highly sampled neonate (z=6.55). The parcellation also performed well against the older, single-band WUNDER dataset (z=3.22). Parcels generated from the primary dataset at height thresholds other than 50% also outperformed chance in external datasets as illustrated in **Supplementary Figure 6**. Parcellation homogeneity remained robust even in external datasets with small sample sizes (**Supplementary Figure 1**).

**Figure 4:**
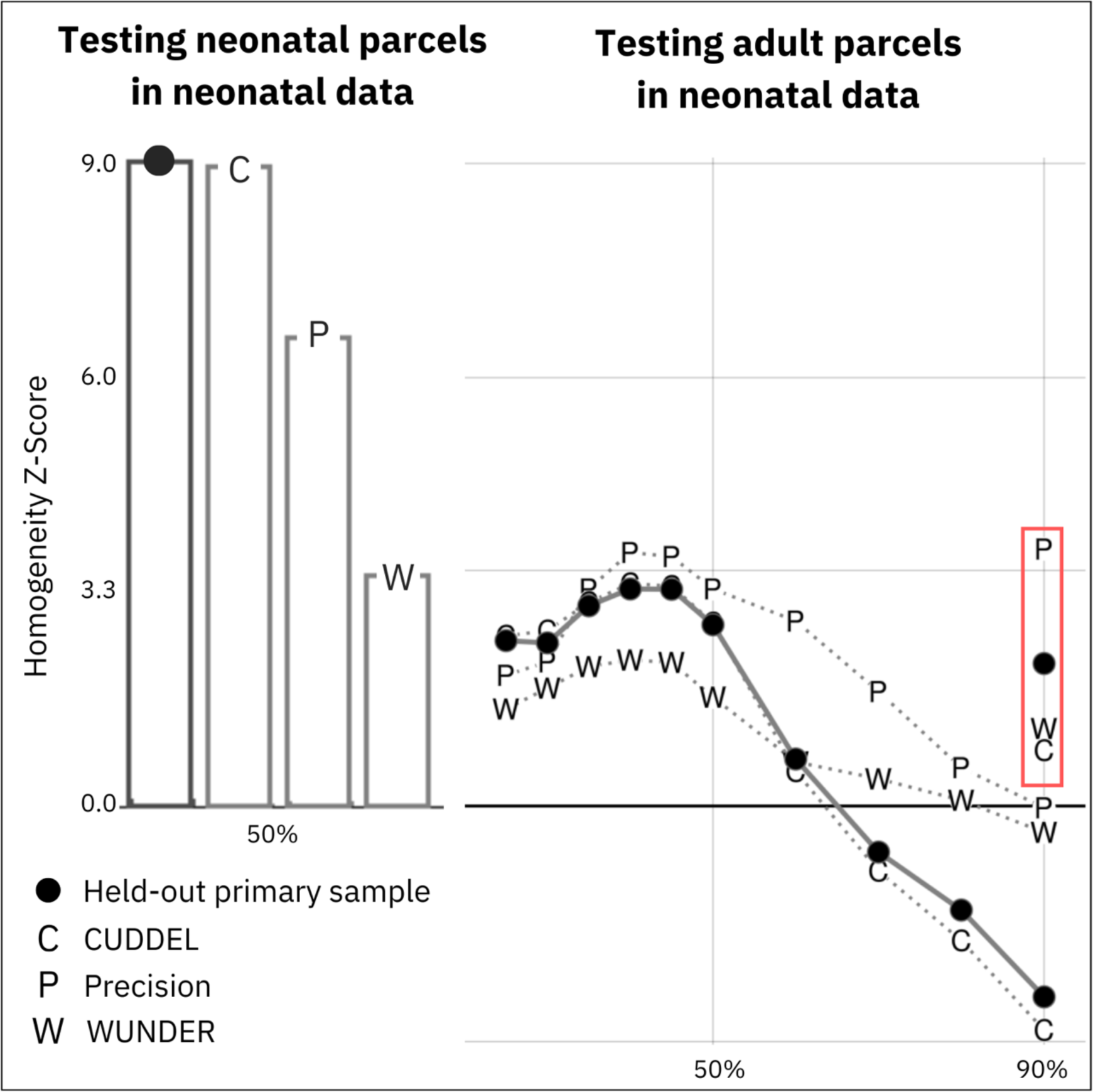
Validation of neonatal parcellations. **A)** The primary dataset (n=131) was used to generate a parcellation which was tested in four datasets. The bar graph represents the homogeneity z-statistic of the parcellation generated from the primary dataset against the second, held-out half of the dataset (n=130; black dot) and three external datasets (CUDDEL+OXYGEN, C; Precision Baby, P; WUNDER, W). **B)** Adult ‘Gordon parcels’ were generated using the same parameters as used in generating the neonatal parcels and were applied to the primary dataset (black dots), as well as external datasets (CUDDEL+OXYGEN, C; Precision Baby, P; WUNDER, W), to test the fit of adult parcels on the neonatal FC data across height thresholds (25%-90%). Markers in the red rectangle denote testing of ‘Gordon parcels’ with original published parameters (90% height threshold; n=400 steps) on each neonatal dataset. Note that adult parcels do a poor job of capturing neonatal FC patterns.

As internal and external validation suggested that the neonatal parcellation performed best using a 50% height criterion, which was substantially different than the criterion previously used for the published adult parcellation (90%; ^6^), we also generated ‘Gordon parcels’ at varying height thresholds (25%-90%) with our modified steps parameter (n=1600). We tested these ‘Gordon parcels’ generated from different height thresholds against our neonatal dataset (Figure 4B), and the adult-based parcels performed no better than chance in the primary neonatal dataset at every height threshold. We further tested the published ‘Gordon parcels’ without any modifications (90% height threshold; n=400 steps) ^6^ on our external neonatal datasets and found that these parcels also performed no better than chance. Thus, the improved fit of neonatal-generated as compared to adult-generated parcels on neonatal data are not due to the difference in the height threshold used to generate the parcels from the boundary map.

### Network Identities of Parcels

We assigned functional network identities to each parcel using the Infomap algorithm. Figure 5 shows ‘consensus’ networks obtained by using information across all edge densities. **Supplementary Figure 7** shows assigned network identities at a selection of edge densities (0.25% to 10%). **Supplementary Table 3** lists all individual parcels by their associated ID number, consensus network assignment, and color in Figure 5. Figure 6 shows a spring-embedded representation of the parcels and their network configuration across 4 representative edge densities. Together, Figures 5 and 6 provide key insights into how neonatal parcels are organized into networks.

**Figure 5:**
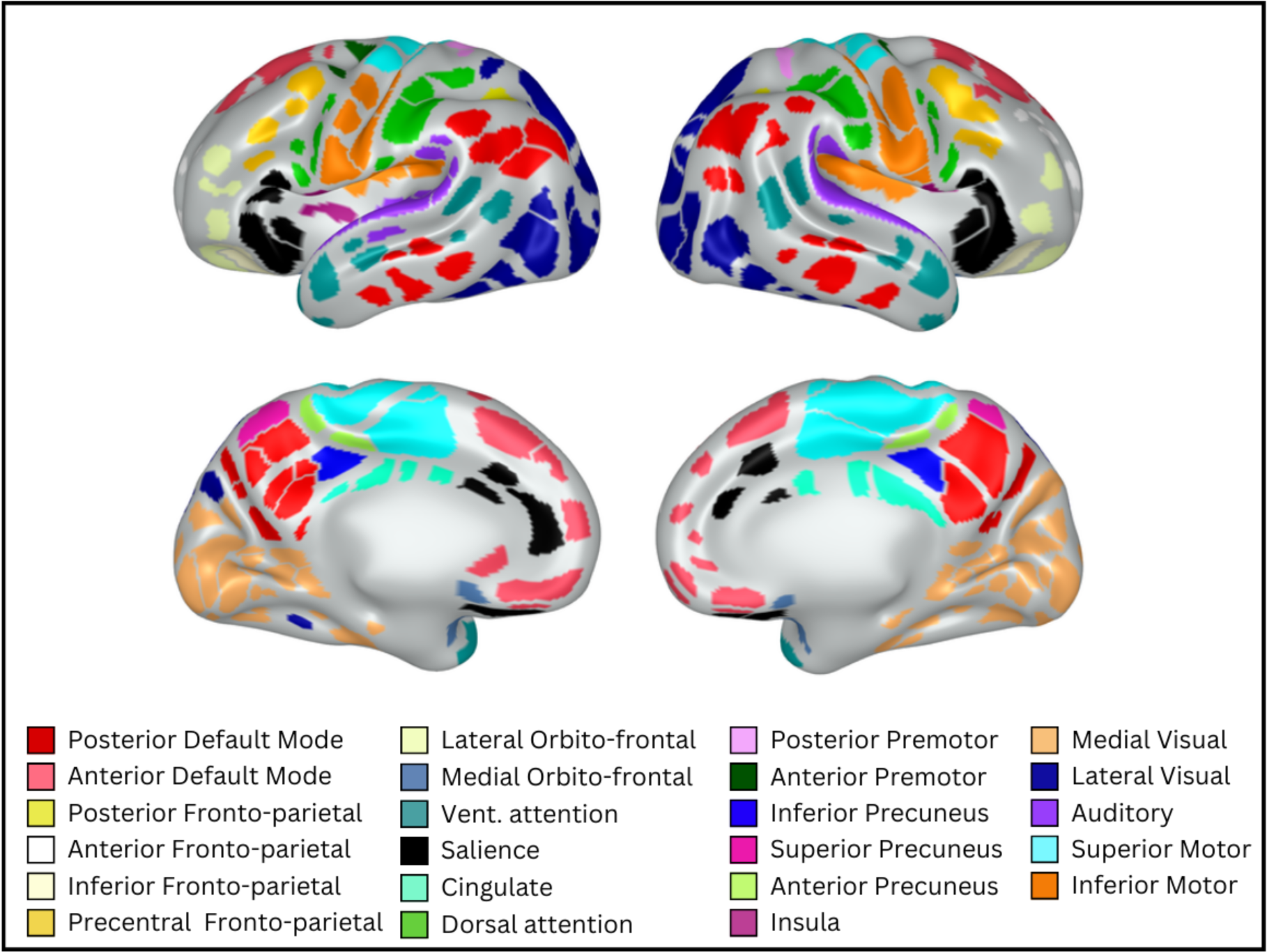
Assigned functional network identities for each parcel. Consensus network assignments for each parcel based on information across all edge densities. Colors and network names were assigned based on adult networks in similar anatomical locations.

**Figure 6:**
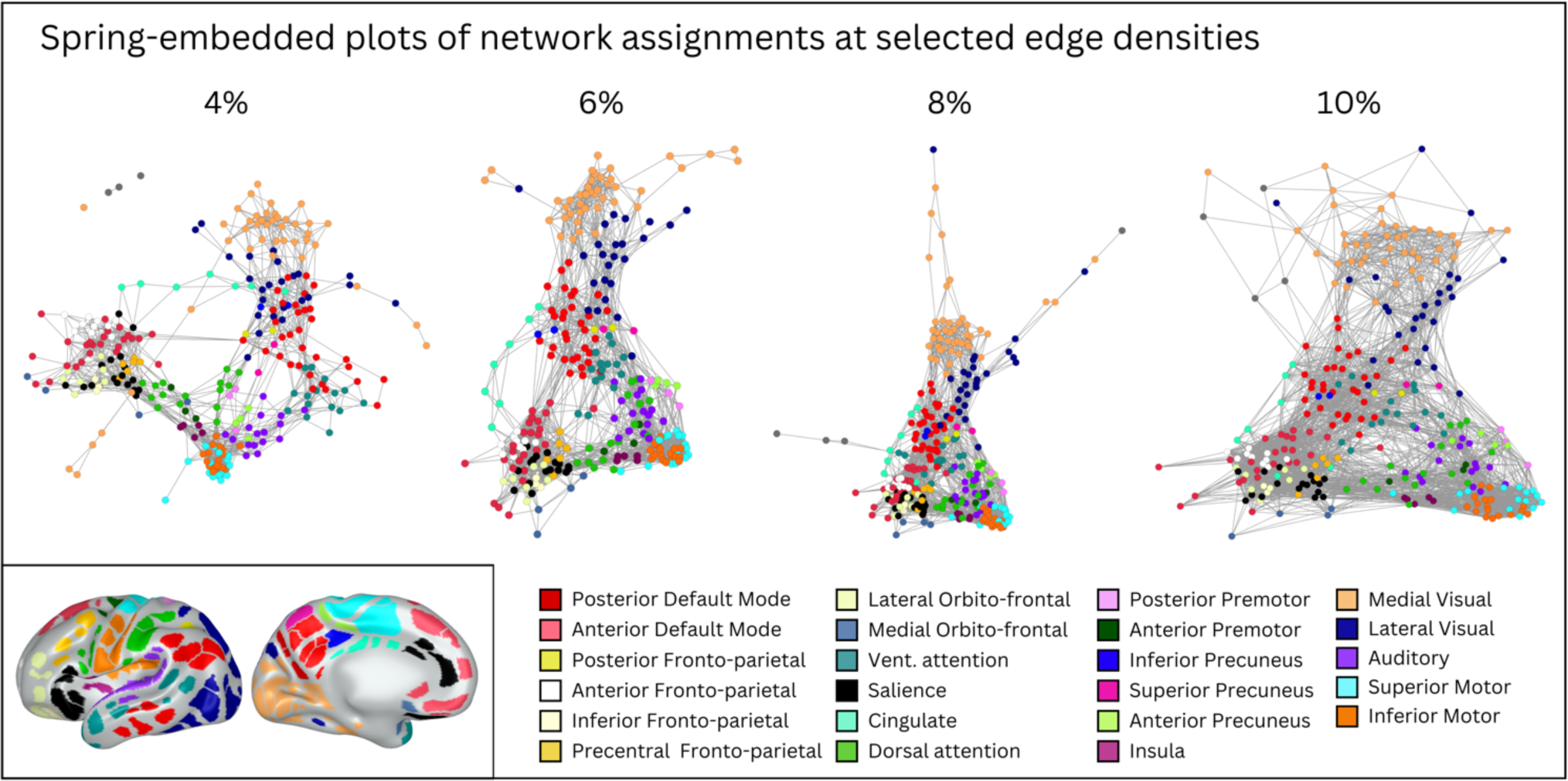
Spring embedded plots of neonatal parcels across different edge densities reveal neonatal network properties. A spring embedded layout of neonatal functional connections at various edge densities between 4% and 10%. Each colored circle corresponds to a particular parcel, colored based on consensus network assignment. Lines represent functional connections between parcels at a given edge density (e.g., at 4% edge density, only the top 4% of positive functional connections are shown). In the spring-embedded representation, stronger connections tend to pull parcels closer together, and so the proximity of parcels to each other is related to their inter-connectivity. Note that when only considering the strongest connections (e.g., 4%), parcels cluster mainly by network (color) and anatomical location (e.g., frontal networks are all near each other); but when considering weaker connections (e.g., 10%), there is some evidence of selective connections between anatomically distant parcels that end up forming the same adult network (e.g., the red and watermelon parcels draw close to each other; these parcels may be precursors of the adult default mode network). Inset shows consensus network assignment of each parcel for reference (identical to Figure 5).

In general, parcels tended to group into networks reminiscent of anatomically isolated sub-portions of adult networks (i.e., most neonatal ‘networks’ only included sets of physically adjacent parcels). For example, the four networks in shades of yellow (see **Supplementary Table 3** for color names) each included only physically adjacent parcels, but together the four networks cover spatially distributed portions of cortex roughly corresponding to the adult fronto-parietal network (FPN). Similarly, the two networks in shades of red together cover portions of cortex roughly corresponding to the adult default mode network (DMN). Additional networks were identified similar to adult dorsal attention (DAN), ventral attention, salience, premotor, motor, visual, and auditory networks. Both the DAN and the posterior DMN included parcels distributed across cortex that were not physically adjacent. Notably, we did not identify a network that clearly corresponded to the cingulo-opercular network (CON); though we did identify several networks that we named based on anatomical location (e.g., cingulate network) that may correspond to sub-portions of the CON.

Spring-embedded representations (Figure 6) are useful for examining within- and between-network relations of the entire parcellation. In such representations, stronger connections tend to pull parcels closer together, and so the proximity of parcels to each other is related to their inter-connectivity. Spring embedded representations can be drawn when considering different edge densities, i.e., considering only the strongest functional connections (low edge densities) or considering progressively weaker functional connections (higher edge densities).

The spring embedded plots reveal that at sparse edge densities that include only the strongest functional connections, parcels cluster by network (color) and then by proximity on the cortical surface (frontal vs. posterior cortex). At the 4% edge density, for example, the frontal lobe networks are in close proximity to each other with few connections to the posterior networks. At denser edge densities, however, (e.g., 10%), there are more functional connections between frontal and posterior parcels, with some selectivity in these connections based on adult network properties. For example, the neonatal networks that putatively correspond to the posterior and anterior portions of the adult-DMN seem to draw together at denser edge densities, as do the posterior and anterior portions of the adult-FPN.

The neonatal connectivity matrix (**Supplementary Figure 8**) also reflected the nascent features of adult-like organization. For example, the motor networks were all positively correlated with each other but negatively correlated with parcels from remaining networks, and the two halves of the putative DMN showed positive correlation with each other.

## DISCUSSION

The current study generated a set of 283 highly homogeneous, reproducible, and externally validated parcels on the cortical surface of the neonatal brain. Parcels from the published literature generated from older infants and adults, in contrast, provided a poor fit to neonatal data. The boundaries denoting transitions in group-average neonatal connectivity between homogeneous parcels were thicker and smoother compared to adult datasets, possibly due to heterogeneity across neonates in the exact placement of parcel borders. As a result, the most generalizable results were obtained when restricting parcels to cover 50% of the neonatal cortical surface. This neonatal parcellation showed high validity both within-sample and across three external datasets, including neonates with a large range of socioeconomic backgrounds and *in utero* drug exposure. Network assignments derived for the neonatal parcels were consistent with prior work: ‘networks’ consisted largely of anatomically adjacent clusters of parcels, but specific sets of neonatal networks together covered anatomical territory reminiscent of adult networks.

### Adult and older infant parcellations do not fit neonatal data

Adult and older-infant brain parcellations generally performed no better than chance in neonatal data, emphasizing the need for an age-specific neonatal parcellation. The neonatal brain is about one-fourth the size of the adult brain, and expansion of the cortical surface over development is non-linear and non-uniform ^16,17^. This non-uniform expansion is the most likely explanation for the poor fit of adult parcels on neonatal data. A consequence is that neuroimaging studies that use adult parcels on neonatal data will not capture functionally homogenous portions of cortex, and thus studies that use adult parcels could provide misleading results. For example, measuring FC of specific ‘Gordon parcels’ in neonates is likely to average this metric across multiple different neonatal cortical areas, providing inaccurate measures of FC and potentially misleading results when describing brain-behavior relations.

### Neonatal boundaries between parcels are thicker and smoother than in adults in group-average data

The zones of transition in FC patterns across the cortical surface, or ‘boundaries’ were thicker and smoother in group-average neonatal datasets compared to adults. Because the boundary smoothness of an *individual* neonate was comparable to that of adults, the most likely explanation of this difference is greater variation across different neonates in exact placement of boundaries. This hypothesis is further supported by the observation that measured border smoothness increased with the number of neonates included in the average. The most likely explanation for increased variability of neonatal boundaries is that the neonatal period is a time of rapid development^33^. As a result, differences in structural development (e.g., cortical folding, surface area) make it more difficult to obtain precise registration of functional data across neonates even with surface registration ^9^. A practical consequence is that parcellations that cover large portions of the neonatal brain (e.g., 90%) include portions of the thick boundaries and thus do not provide as good of a fit to good average data as parcellations that are restricted to the wells between the thick boundaries (e.g., the parcellation generated using the 50% height threshold). While we make both the 90% and 50% height threshold parcellations publicly available, we recommend use of the version generated at 50% to increase generalizability.

### Neonatal parcels can be grouped into functional networks

Functional networks are sets of cortical areas that have high inter-connectivity and have shared functional properties ^3^. Functional networks thus represent an important organizational property of the human brain. In the current study, we used the Infomap algorithm to assign each neonatal parcel to a functional brain network, to aid studies that want to contextualize results by network. Consistent with prior work in neonates, empirically derived ‘networks’ in neonates consisted largely of clusters of anatomically adjacent parcels and sets of these networks together comprised the approximate anatomical locations of anatomically distributed adult-like networks. For example, four of the neonatal networks combined together resembled the adult FPN and two of the neonatal networks combined resembled the adult DMN. Spring-embedded representations suggested that there was weak but selective FC between distinct neonatal networks that putatively combined later in development to be a single network, consistent with prior work ^20,23,24,30,31,53^. An important goal of future work is to track how network organizational properties change over development.

### Neonatal parcels provide an important foundation for studies of human brain development

The neonatal period is a critical stage in neural development that serves as a starting point for postnatal experience-dependent learning ^18^. Complex human behaviors are posited to depend on a tightly coordinated sequence of brain development that extends from the *in-utero* period through at least early adulthood ^54^. Many brain illnesses, including many psychiatric and neurological disorders, are thought to have their origin in very early brain development^55–57^. Thus, characterizing neonatal cortical brain areas and their functional network characteristics is an important goal for systems neuroscience investigations into typical and atypical development. Parcels in the current study may represent cortical areas and thus provide an important starting point for characterizing human brain development.

### Limitations

The present study should be viewed considering its limitations. As noted throughout, biological and methodological challenges may make it more difficult to functionally align groups of neonates compared to older samples using purely anatomical landmarks (i.e., surface-based registration). Consistent with this interpretation, some prior work in older infants has used functional brain properties to improve functional alignment across individuals ^9^. We chose not to incorporate functional alignment in the current study so that the derived parcels would be most generalizable for future neonatal neuroimaging studies, where in most cases it will be impractical to include functional alignment. Both the primary dataset and two of the external datasets were collected on Siemens Prisma 3T scanners, and a third validation dataset was collected on an older Siemens Trio; future work is required to determine how well the generated parcels fits data acquired on other scanners. All neonatal data in the current study were collected during natural sleep, and while the location cortical areas should not depend on sleep state, future work could ascertain how well the parcels fit functional data from awake neonates. Finally, our neonates ranged in age from 38 to 45 weeks PMA, and additional work is required to determine the specific ages in which it is most appropriate to use these parcels.

### Conclusions

We generated a highly reliable set of 283 surface-based parcels for the neonatal brain that were validated using three external datasets. We additionally provide functional brain network assignments for each parcel. This parcellation will aid neonatal neuroimaging studies that seek to describe results contextualized by functionally relevant cortical areas.

## RESOURCE AVAILABILITY

### Lead contact

Further information and requests for resources should be directed to and will be fulfilled by the senior author and lead contact, Chad Sylvester.

### Materials availability

The derived parcellation and the code used to derive the parcels in this work are publicly available at https://github.com/myersm0/myers-labonte_parcellation/. Based on the results above, for typical group studies of neonates, we recommend use of the parcels at 50% height threshold, as depicted in Figures 4 and 5. These parcels are expected to be highly valid across different datasets and individuals, while still covering much of the cortical surface. We also make available the parcels across a range of other height thresholds, including the 90% threshold that covers most of the cortical surface; but note that these parcels are not expected to be as valid across all datasets and subjects. Parcels in the available dataset are numbered according to **Supplementary Table 3**, which also includes network identification. **Supplementary Figure 9** shows each parcel labeled with its parcel number and network identification.

### Data and code availability

● All original code has been deposited at https://github.com/myersm0/WatershedParcellation.jl and is publicly available as of the date of publication.
● The parcellation described in this paper as well as key descriptors (ID, network assignment) are available at https://github.com/myersm0/myers-labonte_parcellation/
● All data reported in this paper will be shared by the lead contact upon request as per study governance procedures.

## ACKNOWLEDGMENTS

This work was supported by funds provided by the Intellectual and Developmental Disabilities Research Center at Washington University P50 HD103525), the Taylor Family Foundation, the National Institute of Mental Health (R01MH122389; R01MH131584, R01MH090786), and the National Institute on Drug Abuse (R01DA046224).

We thank the families of all of our participants from these studies for their participation and dedication towards research.

## AUTHOR CONTRIBUTIONS

Conceptualization, MJM, CMS; Methodology, MJM, EMG, TOL, JCT, DAF, SEP; Software, MJM; Formal Analysis, MJM, AKL, JCT; Resources, NR, JLL, BBW, DMB, CER, CDS, CMS, DAF; Data Curation, MJM; Writing – Original Draft, MJM, AKL, CMS; Writing – Review & Editing, AKL, EMG, TOL, JCT, MDW, ANN, RS, MCC, DMB, CER, CDS, SEP; Visualization, MJM, AKL, JCT; Supervision, CMS; Funding Acquisition, BBW, NR, JLL, DMB, CER, CDS, CMS

