## Supplementary Materials for "Functional parcellation of the neonatal brain"

### SUPPLEMENTARY METHODS

#### Performance of Publicly Available Adult and Infant Parcels on Neonatal Data

To demonstrate how much improvement our neonatal-specific parcellation provides compared to adult parcellations in terms of within-parcel FC homogeneity, we also evaluated our neonatal parcellation created from the primary dataset against several adult surface-based parcellations most commonly used in the neuroimaging literature [1–3], as well as 2 recently published infant parcels developed using datasets including young infants [4–6], on the 130 healthy, term neonates from the neonatal dataset which were not used in the generation of the parcellation.

The Wang parcellations were provided in the HCP standard fsLR 32k surface space by the authors. The Shen parcellation [5,6] was provided in the MNI volume space by the authors and was manually transformed to surface space using the Conte69 midthickness fsLR 32k surfaces using the Connectome Workbench command: "wb_command –volume-label-to-surface-mapping". The resulting file was dilated using "wb_command –cifti-dilate" with 10mm to remove all holes from the volume-to-surface transformation. Next, the isolated patches with a connected component size smaller than 0.1 of the average parcel size of each hemisphere were replaced with the mode of their neighboring vertices using the code from the CBIG repository:
(https://github.com/ThomasYeoLab/CBIG/tree/master/utilities/matlab/parcellation/CBIG_CleanSurfaceParcellation.m). A final step removed parcels smaller than 10 vertices.

Area homogeneity of each parcel was calculated as the proportion of variance across all vertices' connectivity patterns that can be explained by the first principal component [3] and the homogeneity of the parcellation is the average of the homogeneity of all parcels within the parcellation. A homogeneity Z-score was also calculated using the mean and standard deviation of homogeneities from 1000 randomly rotated parcellations on the cortex [3] to account for the potential impact of parcel size and shape on its homogeneity. To create such randomly rotated parcellations, as in analyses in the main text, we rotated each hemisphere of the original parcellation a random amount around each of the x, y and z axes on the spherical expansion of the 32k fs_LR cortical surface. Each parcel was then slightly dilated or contracted to adjust for vertices gained or lost due to the non-uniform vertex density across the surface of the sphere, thus maintaining the same number of vertices within the rotated parcel while approximately maintain the same shape. In any random rotation, some parcels will be rotated into the medial wall (where no data exist) or into high signal dropout areas. In cases where area parcels have fewer than 15 vertices outside the high signal dropout areas (defined as regions where mean BOLD percentage change were 25% below the mode value across all voxels on average throughout the entire scan session, i.e., mean BOLD <750 for BOLD 1000 normalized data) or the medial wall, their homogeneities were not calculated and were imputed by the average homogeneity of all random versions of the parcel in the rest of the 1000 rotations.

### SUPPLEMENTARY TABLES

| **Neonatal Characteristics (n=131)** | ***n*** | **Mean** | **SD** |
|---|---|---|---|
| Sex | | | |
| Male | 78 | | |
| Female | 53 | | |
| Gestational age at birth (weeks) | | 38.5 | 0.99 |
| Postmenstrual age at scan (weeks) | | 41.2 | 1.2 |
| Birthweight in grams | | 3267.9 | 466.1 |
| Area deprivation index | | 69.4 | 24.4 |
| Child's race | | | |
| African American | 48 | | |
| White | 81 | | |
| Chinese | 2 | | |
| Other Pacific Islander | 0 | | |
| Other | 0 | | |
| Mixed African American/White | 0 | | |
| Mixed Chinese/White | 0 | | |
| Ethnicity | | | |
| Hispanic | 3 | | |
| Non-Hispanic | 128 | | |
| **Neonatal fMRI Characteristics** | ***n*** | **Mean** | **SD** |
| fMRI data collected (minutes) | | 22.8 | 2.7 |
| fMRI data retained (minutes) | | 20.2 | 2.8 |

**Supplementary Table 1:** *Demographics for the primary generation dataset which the neonatal boundary map and parcels were generated from.*

| **Neonatal Characteristics (n=66)** | ***n*** | **Mean** | **SD** |
|---|---|---|---|
| Sex | | | |
| Male | 39 | | |
| Female | 27 | | |
| Gestational age at birth (weeks) | | 38.5 | 1.0 |
| Postmenstrual age at scan (weeks) | | 41.2 | 1.3 |
| Birthweight in grams | | 3238.5 | 467.5 |
| Area deprivation index | | 71.8 | 25.1 |
| Child's race | | | |
| African American | 44 | | |
| White | 21 | | |
| Chinese | 1 | | |
| Other Pacific Islander | 0 | | |
| Other | 0 | | |
| Mixed African American/White | 0 | | |
| Mixed Chinese/White | 0 | | |
| Ethnicity | | | |
| Hispanic | 1 | | |
| Non-Hispanic | 65 | | |
| **Neonatal fMRI Characteristics** | ***n*** | **Mean** | **SD** |
| fMRI data collected (minutes) | | 22.7 | 3.0 |
| fMRI data retained (minutes) | | 20.3 | 3.3 |

| **Neonatal Characteristics (n=65)** | ***n*** | **Mean** | **SD** |
|---|---|---|---|
| Sex | | | |
| Male | 39 | | |
| Female | 26 | | |
| Gestational age at birth (weeks) | | 38.5 | 0.97 |
| Postmenstrual age at scan (weeks) | | 41.1 | 1.2 |
| Birthweight in grams | | 3297.6 | 466.4 |
| Area deprivation index | | 70.0 | 23.6 |
| Child's race | | | |
| African American | | 37 | |
| White | | 27 | |
| Chinese | | 1 | |
| Other Pacific Islander | | 0 | |
| Other | | 0 | |
| Mixed African American/White | | 0 | |
| Mixed Chinese/White | | 0 | |
| Ethnicity | | | |
| Hispanic | | 2 | |
| Non-Hispanic | | 63 | |
| **Neonatal fMRI Characteristics** | ***n*** | **Mean** | **SD** |
| fMRI data collected (minutes) | | 23.0 | 2.4 |
| fMRI data retained (minutes) | | 20.1 | 2.1 |

**Supplementary Table 2:** *Demographic information for each split half of the primary dataset.*

| Network Assignment Name | Number Assignment | Color Assignment | RGB Color Code | | |
|---|---|---|---|---|---|
| Unassigned | 0 | Gray | 128 | 128 | 128 |
| Posterior Default Mode | 1 | Red | 255 | 0 | 0 |
| Anterior Default Mode | 2 | Watermelon | 235 | 61 | 84 |
| Superior Precuneus | 3 | Hot Pink | 255 | 26 | 185 |
| Posterior Fronto-parietal | 4 | Yellow | 230 | 230 | 0 |
| Anterior Fronto-parietal | 5 | White | 255 | 255 | 255 |
| Inferior Fronto-parietal | 6 | Cream | 255 | 253 | 208 |
| Precentral Fronto-parietal | 7 | Mustard | 252 | 194 | 0 |
| Lateral Orbito-Frontal | 8 | Citron | 237 | 255 | 176 |
| Medial Orbito-Frontal | 9 | UCLA Blue | 79 | 123 | 176 |
| Salience | 10 | Black | 0 | 0 | 0 |
| Ventral Attention | 11 | Teal | 0 | 153 | 153 |
| Cingulate | 12 | Toothpaste | 0 | 255 | 193 |
| Dorsal Attention | 13 | Green | 0 | 204 | 0 |
| Posterior Premotor | 14 | Pink | 255 | 153 | 255 |
| Anterior Premotor | 15 | Forest Green | 0 | 102 | 0 |
| Inferior Precuneus | 16 | Royal Blue | 0 | 0 | 255 |
| Anterior Precuneus | 17 | Lime | 165 | 255 | 80 |
| Medial Visual | 18 | Tan | 255 | 179 | 102 |
| Lateral Visual | 19 | Dark Blue | 0 | 0 | 153 |
| Auditory | 20 | Purple | 153 | 51 | 255 |
| Insula | 21 | Plum | 148 | 0 | 108 |
| Superior Motor | 22 | Cyan | 0 | 255 | 255 |
| Inferior Motor | 23 | Orange | 255 | 128 | 0 |

| Parcel Number | Network Assignment | Network Assignment Name | Network Assignment Color |
|---|---|---|---|
| 1 | 1 | Posterior Default Mode | Red |
| 2 | 14 | Posterior Premotor | Pink |
| 3 | 2 | Anterior Default Mode | Watermelon |
| 4 | 19 | Lateral Visual | Dark Blue |
| 5 | 1 | Posterior Default Mode | Red |
| 6 | 1 | Posterior Default Mode | Red |
| 7 | 23 | Inferior Motor | Orange |
| 8 | 11 | Ventral Attention | Teal |
| 9 | 12 | Cingulate | Toothpaste |
| 10 | 18 | Medial Visual | Tan |

| | | | |
|---|---|---|---|
| 11 | 18 | Medial Visual | Tan |
| 12 | 18 | Medial Visual | Tan |
| 13 | 1 | Posterior Default Mode | Red |
| 14 | 18 | Medial Visual | Tan |
| 15 | 22 | Superior Motor | Cyan |
| 16 | 10 | Salience | Black |
| 17 | 10 | Salience | Black |
| 18 | 12 | Cingulate | Toothpaste |
| 19 | 12 | Cingulate | Toothpaste |
| 20 | 12 | Cingulate | Toothpaste |
| 21 | 10 | Salience | Black |
| 22 | 1 | Posterior Default Mode | Red |
| 23 | 1 | Posterior Default Mode | Red |
| 24 | 22 | Superior Motor | Cyan |
| 25 | 22 | Superior Motor | Cyan |
| 26 | 22 | Superior Motor | Cyan |
| 27 | 17 | Anterior Precuneus | Lime |
| 28 | 16 | Inferior Precuneus | Royal Blue |
| 29 | 17 | Anterior Precuneus | Lime |
| 30 | 23 | Inferior Motor | Orange |
| 31 | 23 | Inferior Motor | Orange |
| 32 | 13 | Dorsal Attention | Green |
| 33 | 2 | Anterior Default Mode | Watermelon |
| 34 | 2 | Anterior Default Mode | Watermelon |
| 35 | 22 | Superior Motor | Cyan |
| 36 | 22 | Superior Motor | Cyan |
| 37 | 15 | Anterior Premotor | Forest Green |
| 38 | 15 | Anterior Premotor | Forest Green |
| 39 | 2 | Anterior Default Mode | Watermelon |
| 40 | 14 | Posterior Premotor | Pink |
| 41 | 13 | Dorsal Attention | Green |
| 42 | 23 | Inferior Motor | Orange |
| 43 | 13 | Dorsal Attention | Green |
| 44 | 13 | Dorsal Attention | Green |
| 45 | 22 | Superior Motor | Cyan |
| 46 | 23 | Inferior Motor | Orange |
| 47 | 23 | Inferior Motor | Orange |
| 48 | 1 | Posterior Default Mode | Red |
| 49 | 11 | Ventral Attention | Teal |
| 50 | 20 | Auditory | Purple |
| 51 | 20 | Auditory | Purple |

| | | | |
|---|---|---|---|
| 52 | 23 | Inferior Motor | Orange |
| 53 | 1 | Posterior Default Mode | Red |
| 54 | 20 | Auditory | Purple |
| 55 | 20 | Auditory | Purple |
| 56 | 20 | Auditory | Purple |
| 57 | 11 | Ventral Attention | Teal |
| 58 | 11 | Ventral Attention | Teal |
| 59 | 20 | Auditory | Purple |
| 60 | 20 | Auditory | Purple |
| 61 | 20 | Auditory | Purple |
| 62 | 20 | Auditory | Purple |
| 63 | 20 | Auditory | Purple |
| 64 | 8 | Lateral Orbito-Frontal | Citron |
| 65 | 10 | Salience | Black |
| 66 | 21 | Insula | Plum |
| 67 | 6 | Inferior Fronto-parietal | Cream |
| 68 | 6 | Inferior Fronto-parietal | Cream |
| 69 | 9 | Medial Orbito-Frontal | UCLA Blue |
| 70 | 21 | Insula | Plum |
| 71 | 10 | Salience | Black |
| 72 | 10 | Salience | Black |
| 73 | 8 | Lateral Orbito-Frontal | Citron |
| 74 | 19 | Lateral Visual | Dark Blue |
| 75 | 1 | Posterior Default Mode | Red |
| 76 | 1 | Posterior Default Mode | Red |
| 77 | 18 | Medial Visual | Tan |
| 78 | 18 | Medial Visual | Tan |
| 79 | 3 | Superior Precuneus | Hot Pink |
| 80 | 19 | Lateral Visual | Dark Blue |
| 81 | 1 | Posterior Default Mode | Red |
| 82 | 1 | Posterior Default Mode | Red |
| 83 | 1 | Posterior Default Mode | Red |
| 84 | 19 | Lateral Visual | Dark Blue |
| 85 | 1 | Posterior Default Mode | Red |
| 86 | 19 | Lateral Visual | Dark Blue |
| 87 | 4 | Posterior Fronto-parietal | Yellow |
| 88 | 19 | Lateral Visual | Dark Blue |
| 89 | 19 | Lateral Visual | Dark Blue |
| 90 | 19 | Lateral Visual | Dark Blue |
| 91 | 19 | Lateral Visual | Dark Blue |
| 92 | 19 | Lateral Visual | Dark Blue |

| | | | |
|---|---|---|---|
| 93 | 19 | Lateral Visual | Dark Blue |
| 94 | 1 | Posterior Default Mode | Red |
| 95 | 23 | Inferior Motor | Orange |
| 96 | 23 | Inferior Motor | Orange |
| 97 | 13 | Dorsal Attention | Green |
| 98 | 7 | Precentral Fronto-parietal | Mustard |
| 99 | 8 | Lateral Orbito-Frontal | Citron |
| 100 | 10 | Salience | Black |
| 101 | 7 | Precentral Fronto-parietal | Mustard |
| 102 | 21 | Insula | Plum |
| 103 | 13 | Dorsal Attention | Green |
| 104 | 13 | Dorsal Attention | Green |
| 105 | 13 | Dorsal Attention | Green |
| 106 | 5 | Anterior Fronto-parietal | White |
| 107 | 2 | Anterior Default Mode | Watermelon |
| 108 | 9 | Medial Orbito-Frontal | UCLA Blue |
| 109 | 8 | Lateral Orbito-Frontal | Citron |
| 110 | 10 | Salience | Black |
| 111 | 2 | Anterior Default Mode | Watermelon |
| 112 | 2 | Anterior Default Mode | Watermelon |
| 113 | 1 | Posterior Default Mode | Red |
| 114 | 1 | Posterior Default Mode | Red |
| 115 | 11 | Ventral Attention | Teal |
| 116 | 18 | Medial Visual | Tan |
| 117 | 18 | Medial Visual | Tan |
| 118 | 0 | Unassigned | Gray |
| 119 | 18 | Medial Visual | Tan |
| 120 | 19 | Lateral Visual | Dark Blue |
| 121 | 19 | Lateral Visual | Dark Blue |
| 122 | 19 | Lateral Visual | Dark Blue |
| 123 | 19 | Lateral Visual | Dark Blue |
| 124 | 18 | Medial Visual | Tan |
| 125 | 18 | Medial Visual | Tan |
| 126 | 18 | Medial Visual | Tan |
| 127 | 18 | Medial Visual | Tan |
| 128 | 18 | Medial Visual | Tan |
| 129 | 18 | Medial Visual | Tan |
| 130 | 18 | Medial Visual | Tan |
| 131 | 18 | Medial Visual | Tan |
| 132 | 0 | Unassigned | Gray |
| 133 | 18 | Medial Visual | Tan |

| | | | |
|---|---|---|---|
| 134 | 5 | Anterior Fronto-parietal | White |
| 135 | 2 | Anterior Default Mode | Watermelon |
| 136 | 5 | Anterior Fronto-parietal | White |
| 137 | 2 | Anterior Default Mode | Watermelon |
| 138 | 2 | Anterior Default Mode | Watermelon |
| 139 | 2 | Anterior Default Mode | Watermelon |
| 140 | 7 | Precentral Fronto-parietal | Mustard |
| 141 | 7 | Precentral Fronto-parietal | Mustard |
| 142 | 20 | Auditory | Purple |
| 143 | 11 | Ventral Attention | Teal |
| 144 | 11 | Ventral Attention | Teal |
| 145 | 11 | Ventral Attention | Teal |
| 146 | 1 | Posterior Default Mode | Red |
| 147 | 1 | Posterior Default Mode | Red |
| 148 | 22 | Superior Motor | Cyan |
| 149 | 23 | Inferior Motor | Orange |
| 150 | 2 | Anterior Default Mode | Watermelon |
| 151 | 19 | Lateral Visual | Dark Blue |
| 152 | 1 | Posterior Default Mode | Red |
| 153 | 1 | Posterior Default Mode | Red |
| 154 | 23 | Inferior Motor | Orange |
| 155 | 12 | Cingulate | Toothpaste |
| 156 | 18 | Medial Visual | Tan |
| 157 | 18 | Medial Visual | Tan |
| 158 | 18 | Medial Visual | Tan |
| 159 | 18 | Medial Visual | Tan |
| 160 | 18 | Medial Visual | Tan |
| 161 | 18 | Medial Visual | Tan |
| 162 | 22 | Superior Motor | Cyan |
| 163 | 10 | Salience | Black |
| 164 | 10 | Salience | Black |
| 165 | 12 | Cingulate | Toothpaste |
| 166 | 12 | Cingulate | Toothpaste |
| 167 | 10 | Salience | Black |
| 168 | 1 | Posterior Default Mode | Red |
| 169 | 22 | Superior Motor | Cyan |
| 170 | 17 | Anterior Precuneus | Lime |
| 171 | 16 | Inferior Precuneus | Royal Blue |
| 172 | 17 | Anterior Precuneus | Lime |
| 173 | 22 | Superior Motor | Cyan |
| 174 | 22 | Superior Motor | Cyan |

| | | | |
|---|---|---|---|
| 175 | 2 | Anterior Default Mode | Watermelon |
| 176 | 22 | Superior Motor | Cyan |
| 177 | 22 | Superior Motor | Cyan |
| 178 | 15 | Anterior Premotor | Forest Green |
| 179 | 22 | Superior Motor | Cyan |
| 180 | 14 | Posterior Premotor | Pink |
| 181 | 13 | Dorsal Attention | Green |
| 182 | 13 | Dorsal Attention | Green |
| 183 | 23 | Inferior Motor | Orange |
| 184 | 23 | Inferior Motor | Orange |
| 185 | 1 | Posterior Default Mode | Red |
| 186 | 11 | Ventral Attention | Teal |
| 187 | 20 | Auditory | Purple |
| 188 | 20 | Auditory | Purple |
| 189 | 1 | Posterior Default Mode | Red |
| 190 | 1 | Posterior Default Mode | Red |
| 191 | 20 | Auditory | Purple |
| 192 | 11 | Ventral Attention | Teal |
| 193 | 11 | Ventral Attention | Teal |
| 194 | 20 | Auditory | Purple |
| 195 | 11 | Ventral Attention | Teal |
| 196 | 8 | Lateral Orbito-Frontal | Citron |
| 197 | 10 | Salience | Black |
| 198 | 21 | Insula | Plum |
| 199 | 6 | Inferior Fronto-parietal | Cream |
| 200 | 9 | Medial Orbito-Frontal | UCLA Blue |
| 201 | 10 | Salience | Black |
| 202 | 10 | Salience | Black |
| 203 | 10 | Salience | Black |
| 204 | 8 | Lateral Orbito-Frontal | Citron |
| 205 | 19 | Lateral Visual | Dark Blue |
| 206 | 19 | Lateral Visual | Dark Blue |
| 207 | 1 | Posterior Default Mode | Red |
| 208 | 18 | Medial Visual | Tan |
| 209 | 18 | Medial Visual | Tan |
| 210 | 3 | Superior Precuneus | Hot Pink |
| 211 | 1 | Posterior Default Mode | Red |
| 212 | 1 | Posterior Default Mode | Red |
| 213 | 19 | Lateral Visual | Dark Blue |
| 214 | 4 | Posterior Fronto-parietal | Yellow |
| 215 | 19 | Lateral Visual | Dark Blue |

| | | | |
|---|---|---|---|
| 216 | 19 | Lateral Visual | Dark Blue |
| 217 | 19 | Lateral Visual | Dark Blue |
| 218 | 1 | Posterior Default Mode | Red |
| 219 | 1 | Posterior Default Mode | Red |
| 220 | 23 | Inferior Motor | Orange |
| 221 | 23 | Inferior Motor | Orange |
| 222 | 23 | Inferior Motor | Orange |
| 223 | 13 | Dorsal Attention | Green |
| 224 | 23 | Inferior Motor | Orange |
| 225 | 13 | Dorsal Attention | Green |
| 226 | 7 | Precentral Fronto-parietal | Mustard |
| 227 | 7 | Precentral Fronto-parietal | Mustard |
| 228 | 21 | Insula | Plum |
| 229 | 13 | Dorsal Attention | Green |
| 230 | 13 | Dorsal Attention | Green |
| 231 | 13 | Dorsal Attention | Green |
| 232 | 5 | Anterior Fronto-parietal | White |
| 233 | 2 | Anterior Default Mode | Watermelon |
| 234 | 9 | Medial Orbito-Frontal | UCLA Blue |
| 235 | 8 | Lateral Orbito-Frontal | Citron |
| 236 | 5 | Anterior Fronto-parietal | White |
| 237 | 10 | Salience | Black |
| 238 | 10 | Salience | Black |
| 239 | 2 | Anterior Default Mode | Watermelon |
| 240 | 2 | Anterior Default Mode | Watermelon |
| 241 | 2 | Anterior Default Mode | Watermelon |
| 242 | 1 | Posterior Default Mode | Red |
| 243 | 1 | Posterior Default Mode | Red |
| 244 | 11 | Ventral Attention | Teal |
| 245 | 18 | Medial Visual | Tan |
| 246 | 18 | Medial Visual | Tan |
| 247 | 18 | Medial Visual | Tan |
| 248 | 0 | Unassigned | Gray |
| 249 | 18 | Medial Visual | Tan |
| 250 | 19 | Lateral Visual | Dark Blue |
| 251 | 19 | Lateral Visual | Dark Blue |
| 252 | 19 | Lateral Visual | Dark Blue |
| 253 | 19 | Lateral Visual | Dark Blue |
| 254 | 18 | Medial Visual | Tan |
| 255 | 18 | Medial Visual | Tan |
| 256 | 18 | Medial Visual | Tan |

| | | | |
|---|---|---|---|
| 257 | 18 | Medial Visual | Tan |
| 258 | 18 | Medial Visual | Tan |
| 259 | 18 | Medial Visual | Tan |
| 260 | 18 | Medial Visual | Tan |
| 261 | 18 | Medial Visual | Tan |
| 262 | 18 | Medial Visual | Tan |
| 263 | 18 | Medial Visual | Tan |
| 264 | 18 | Medial Visual | Tan |
| 265 | 18 | Medial Visual | Tan |
| 266 | 0 | Unassigned | Gray |
| 267 | 18 | Medial Visual | Tan |
| 268 | 2 | Anterior Default Mode | Watermelon |
| 269 | 5 | Anterior Fronto-parietal | White |
| 270 | 2 | Anterior Default Mode | Watermelon |
| 271 | 5 | Anterior Fronto-parietal | White |
| 272 | 2 | Anterior Default Mode | Watermelon |
| 273 | 5 | Anterior Fronto-parietal | White |
| 274 | 2 | Anterior Default Mode | Watermelon |
| 275 | 2 | Anterior Default Mode | Watermelon |
| 276 | 2 | Anterior Default Mode | Watermelon |
| 277 | 5 | Anterior Fronto-parietal | White |
| 278 | 5 | Anterior Fronto-parietal | White |
| 279 | 2 | Anterior Default Mode | Watermelon |
| 280 | 2 | Anterior Default Mode | Watermelon |
| 281 | 20 | Auditory | Purple |
| 282 | 11 | Ventral Attention | Teal |
| 283 | 11 | Ventral Attention | Teal |

**Supplementary Table 3:** *Network identity table for each parcel.* The top table includes the name of each functional network with its corresponding network number, color, and RBG color codes. In the bottom table, each row contains information about a single parcel's network identity (network number, name, and color).

**SUPPLEMENTARY FIGURES**

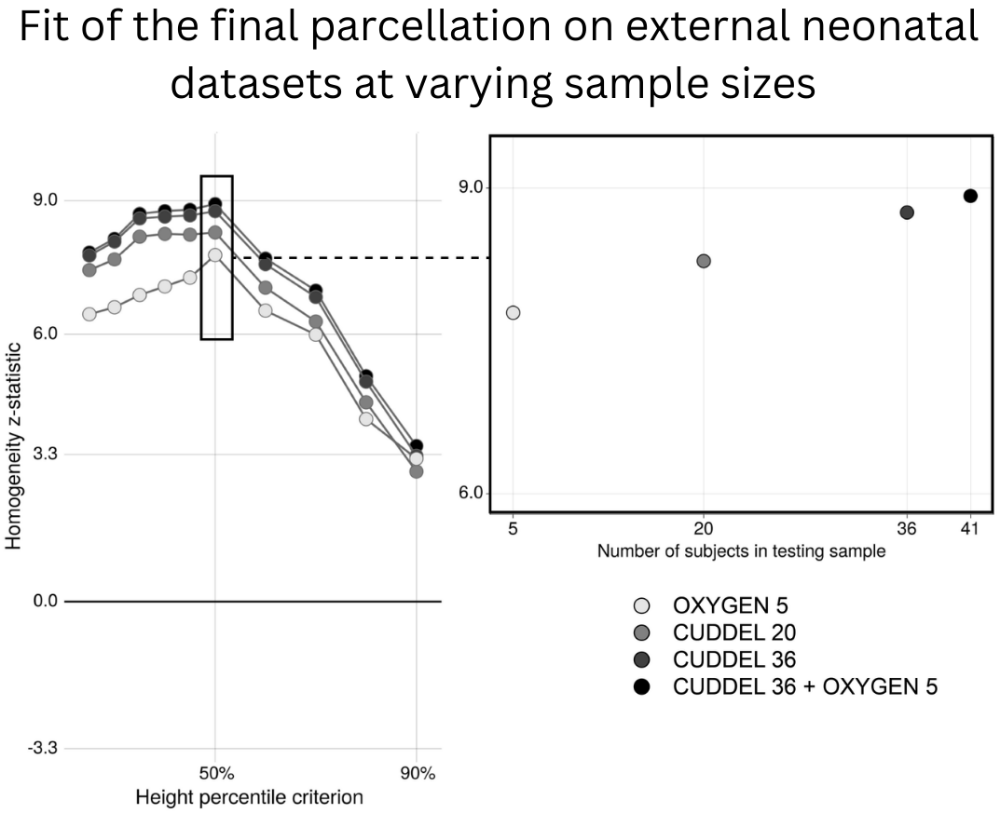


**Supplementary Figure 1:** *Parcellation homogeneity remained robust even with small sample sizes in external validation datasets, CUDDEL and OXYGEN.* **Left:** Homogeneity z-statistic of parcellations generated using the primary dataset (n=131) at different height percentiles tested using data from two external datasets, CUDDEL and OXYGEN. **Right**: Homogeneity z-statistic for the final parcellation (parcels from the 50% height threshold) plotted as a function of sample size for external validation datasets. Color of the dot represents the dataset and the number of subjects included in the analyses.

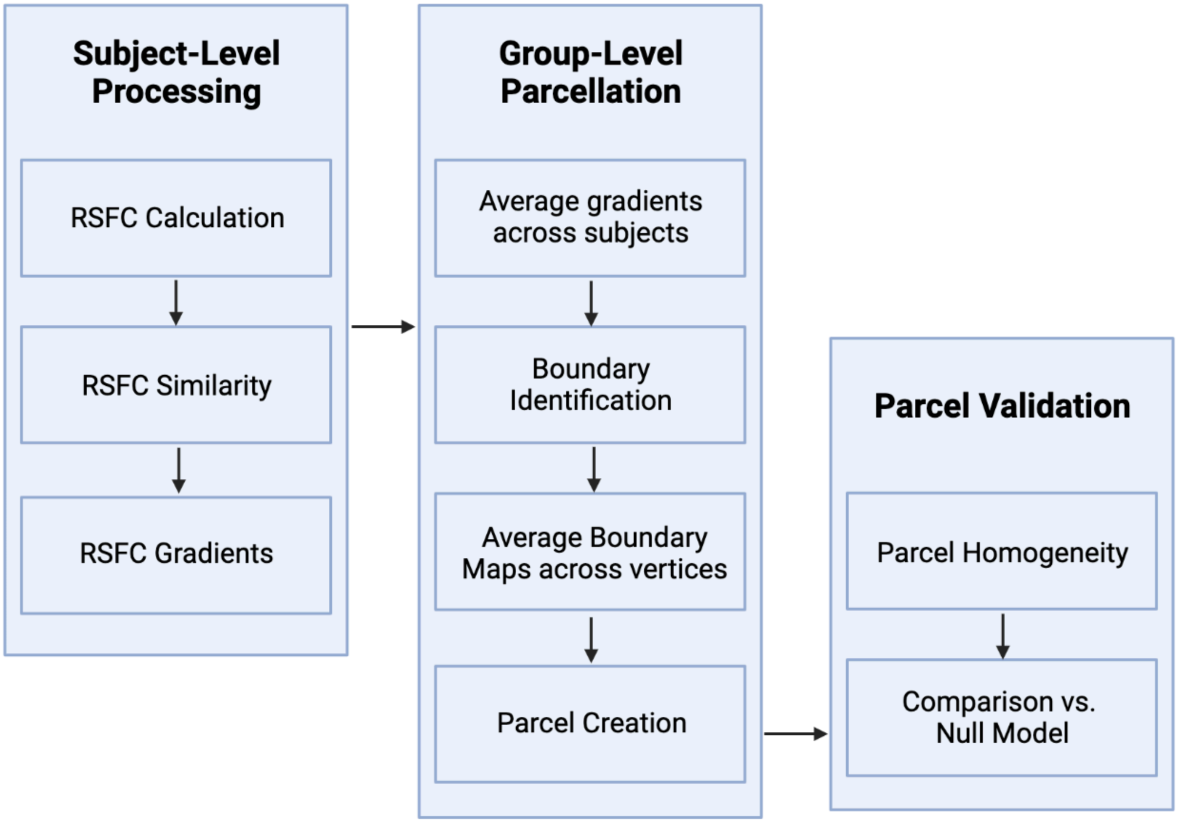


**Supplementary Figure 2:** *Flow of parcellation generation and validation methods.*

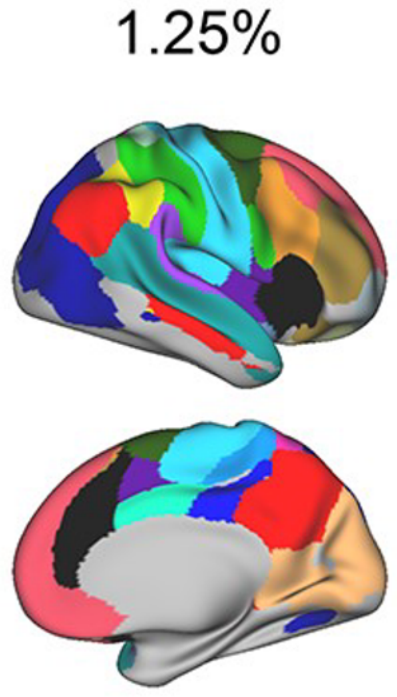


**Supplementary Figure 3:** *Previously described neonate-specific vertex-wise networks from Sylvester et al., 2022 used as a template for generating the consensus network assignments for the generated parcellation.* The network assignments as adapted from Figure 3A from Sylvester et al., 2022 were generated using the standard Infomap approach at the 1.25% edge density[7].

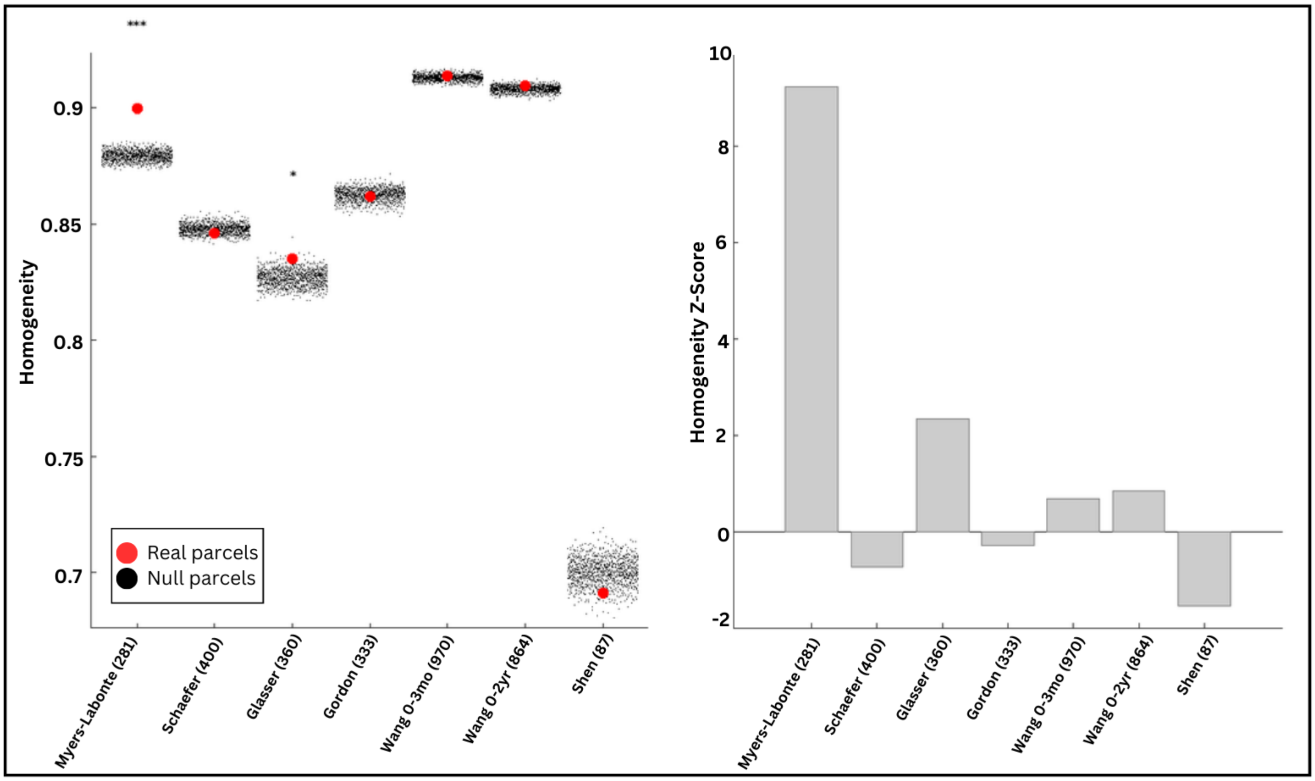


**Supplementary Figure 4:** *Publicly available adult parcellations perform poorly on neonatal data.* Parcellations tested include the Schaefer [2], Glasser [1], and Gordon [3] adult parcellations as well as the Wang [4] and Shen [6] parcellations from older infants. The Myers-Labonte parcellation is the parcellation from the current study. The left panel indicates true averaged homogeneity values of each parcellation tested against neonatal data (red dots) as well as null model homogeneity values for each parcellation generated by rotating the parcellation to random locations on the cortical surface (black dots). The right panel indicates homogeneity z-scores of each parcellation tested against neonatal data. The Glasser parcellation was the only parcellation to perform better than chance in the neonatal data, and its z-score (2.3) was much lower than the z-score of the parcellation generated from neonates in the current study (9.2). Note that the Myers-Labonte parcellation in the current dataset was tested against held-out data not used to generate the parcellation. Parenthetical numbers next to each parcellation name indicate the number of parcels in that parcellation.

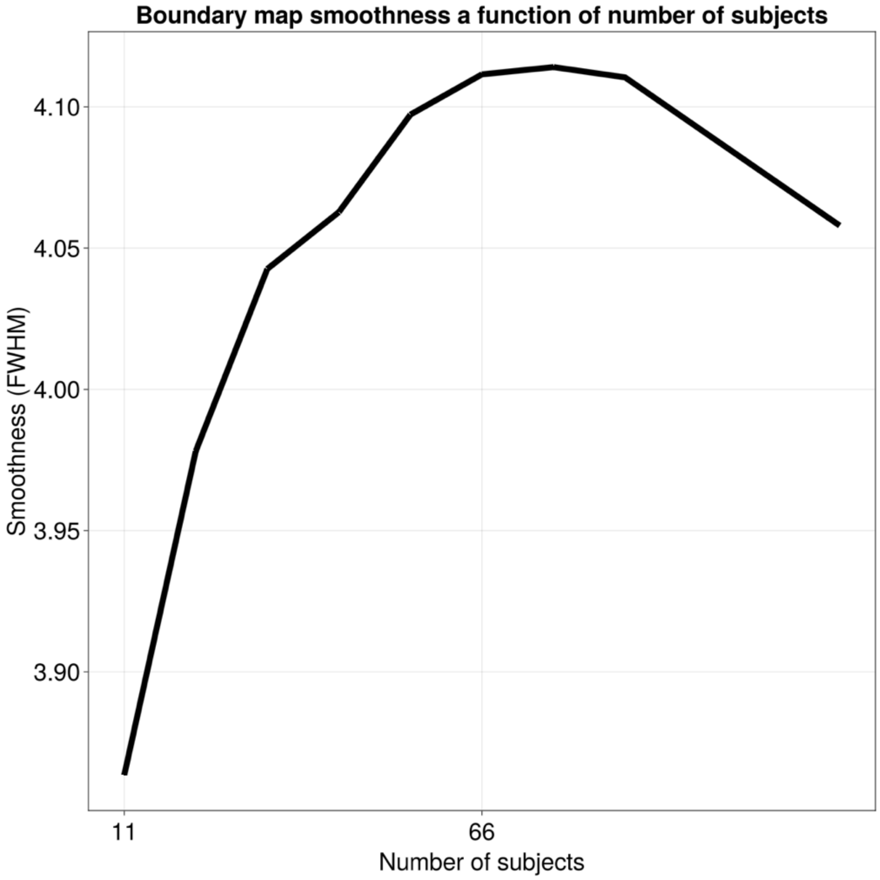


**Supplementary Figure 5:** *Smoothness of the neonatal boundary map increased with increasing number of subjects included in creating the average.*

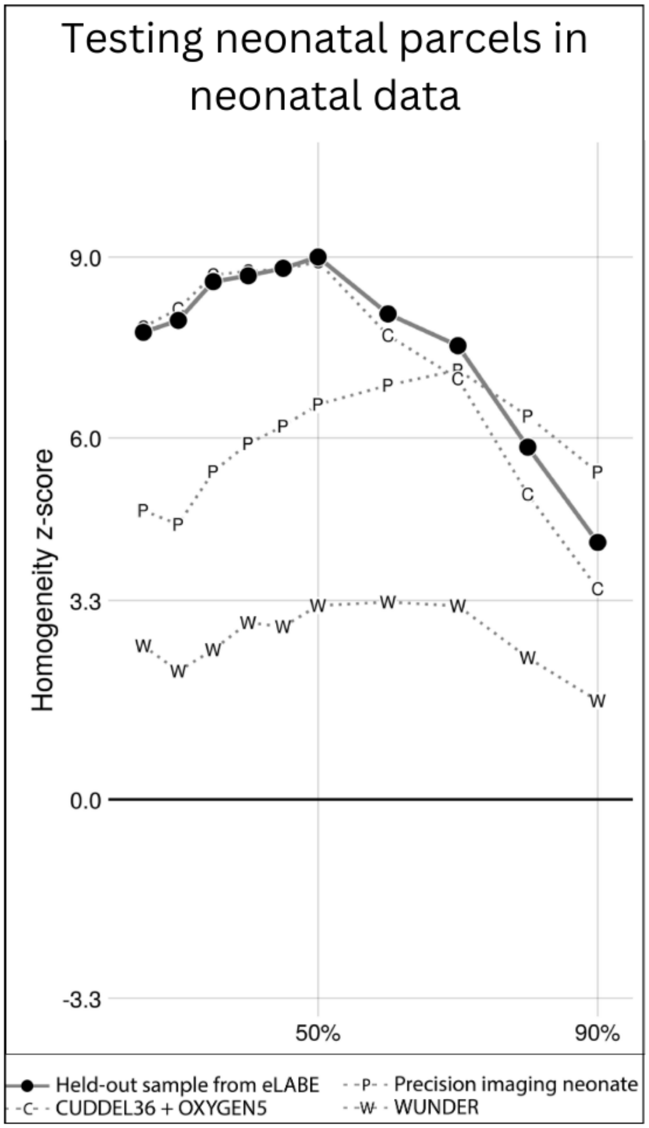


**Supplementary Figure 6:** *Validation of neonatal parcellations.* The primary dataset (n=131) was used to generate parcellations at varying height thresholds between 25% and 90%, which were tested in four datasets. The curves represent the homogeneity z-statistic of the parcellation generated from the primary dataset against the held-out second half of the dataset (n=130; black dots), and three external datasets (CUDDEL+OXYGEN, C; Precision Baby, PB; WUNDER, W). Note that in all cases, parcellations at 50% edge density and lower performed very well when tested against internal and external datasets.

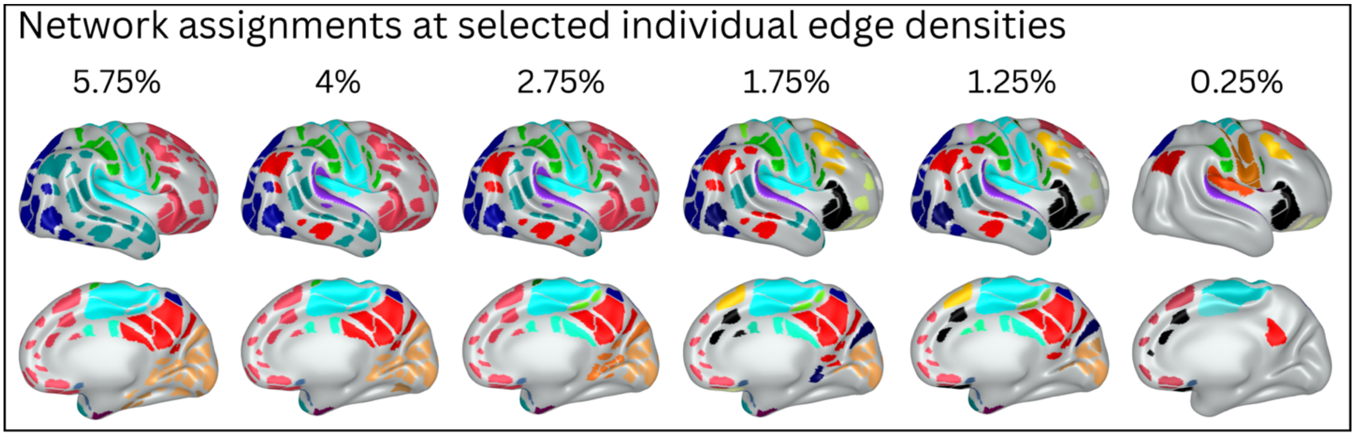


**Supplementary Figure 7:** *Assigned network identities at selected individual edge densities from 0.25% to 5.75%.*

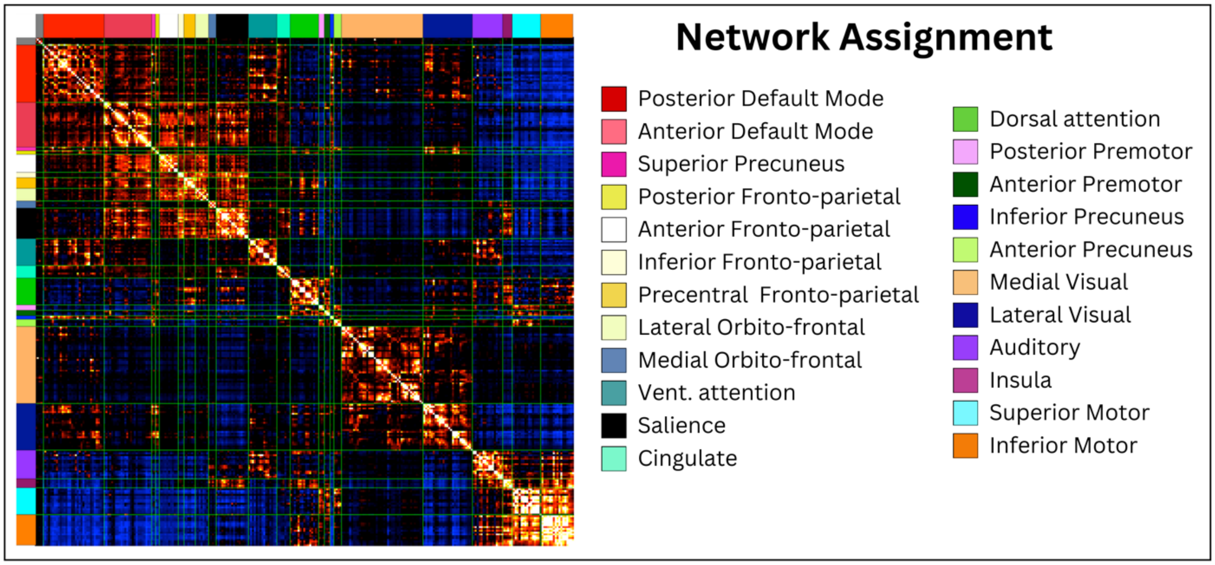


**Supplementary Figure 8:** *Functional connectivity matrix of the generated parcellation with parcels ordered by network identity from the consensus map.*

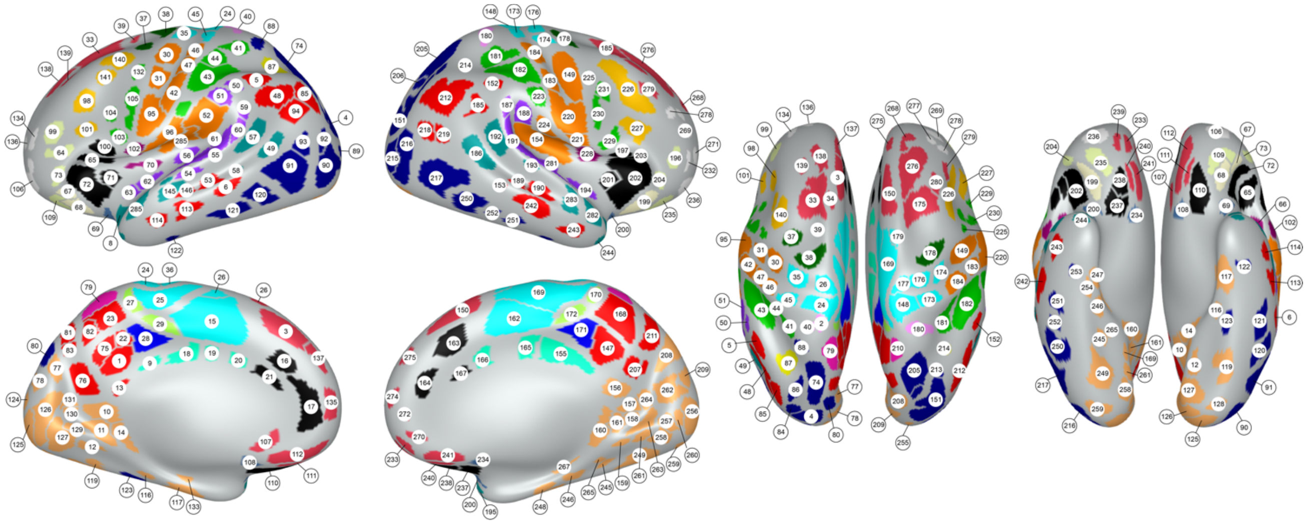

**Supplementary Figure 9:** *Parcels labeled with their respective parcel number and network assignment.*
